# Unlocking the Role of sMyBP-C: A Key Player in Skeletal Muscle Development and Growth

**DOI:** 10.1101/2023.10.23.563591

**Authors:** Taejeong Song, James W. McNamara, Akhil Baby, Weikang Ma, Maicon Landim-Vieira, Sankar Natesan, Jose Renato Pinto, John N. Lorenz, Thomas C. Irving, Sakthivel Sadayappan

## Abstract

Skeletal muscle is the largest organ in the body, responsible for gross movement and metabolic regulation. Recently, variants in the *MYBPC1* gene have been implicated in a variety of developmental muscle diseases, such as distal arthrogryposis. How *MYBPC1* variants cause disease is not well understood. Here, through a collection of novel gene-edited mouse models, we define a critical role for slow myosin binding protein-C (sMyBP-C), encoded by *MYBPC1*, across muscle development, growth, and maintenance during prenatal, perinatal, postnatal and adult stages. Specifically, *Mybpc1* knockout mice exhibited early postnatal lethality and impaired skeletal muscle formation and structure, skeletal deformity, and respiratory failure. Moreover, a conditional knockout of *Mybpc1* in perinatal, postnatal and adult stages demonstrates impaired postnatal muscle growth and function secondary to disrupted actomyosin interaction and sarcomere structural integrity. These findings confirm the essential role of sMyBP-C in skeletal muscle and reveal specific functions in both prenatal embryonic musculoskeletal development and postnatal muscle growth and function.

---

The sarcomere is the fundamental functional unit of skeletal muscle. During muscle development, multiple structural and regulatory sarcomere proteins in thick and thin filaments are expressed spatiotemporally to confer physiological specification to muscle groups. Dysregulation of sarcomere proteins can lead to severe inherited myopathies resulting in immature muscle structure and functional deficits as well as embryonic or postnatal death (1). Myosin binding protein-C is a crucial sarcomere regulatory protein localized in the C-zone of the A band associated with the thick filament (2). Of the three MyBP-C isoforms, slow and fast MyBP-C (sMyBP-C and fMyBP-C) are expressed in skeletal muscle and encoded by *MYBPC1* and *MYBPC2* genes, respectively (Figure 1A). The third MyBP-C isoform, cardiac MyBP-C (cMyBP-C), encoded by *MYBPC3* gene, is specific to the heart (3). cMyBP-C regulates actomyosin interaction and calcium transients in the heart, and mutations in it are closely linked to the development of hypertrophic cardiomyopathy (HCM) (4–6). sMyBP-C is expressed in all muscle fiber types (7). Multiple splice variants and post-translational modifications of sMyBP-C have been identified in human and mouse skeletal muscles (8, 9). The recently identified association of mutations in *MYBPC1* gene with distal arthrogryposis (DA) has increased its clinical relevance (10, 11). While previous studies have reported the occurrence of muscle atrophy and functional deficits in mouse and zebrafish models (12, 13) and lethal congenital contracture syndrome-4 in patients (*14*) lacking sMyBP-C, our understanding of the precise mechanisms through which it regulates muscle structure and function remains incomplete.

**Figure 1.**
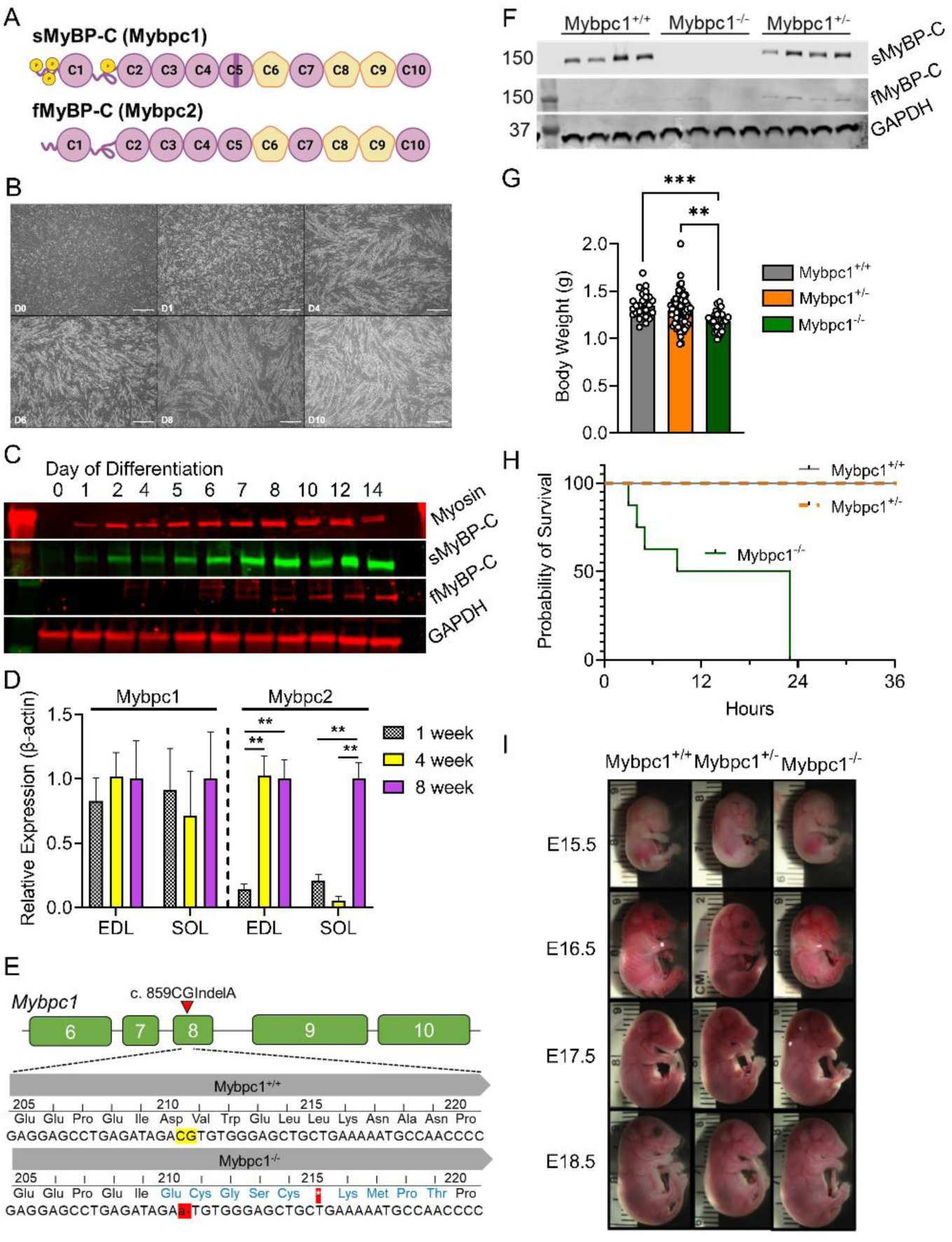
Essential early constitutive sMyBP-C expression for neonatal mouse survival. A) Domain structure of two skeletal MyBP-C isoforms, slow and fast MyBP-C encoded in *Mybpc1* and *Mybpc2* genes, respectively. B) Time course images of differentiating C2C12. Scale bar=500um. C) Early sMyBP-C expression but late expression of fMyBP-C protein in differentiating C2C12 myocytes. D) Expression profile of Mybpc1 and *Mybpc2* genes in mouse fast (EDL) and slow (soleus) twitch muscles at 1, 4 and 8 weeks old (n=3 per group). E) Schematic illustration of a mouse model with sequence comparison of *Mybpc1*^+/+^ and *Mybpc1*^-/-^. F) sMyBP-C and fMyBP-C protein expression in diaphragm muscles (n=4 per group). G) Average body weight measured immediately after birth. H) Survival curve of *Mybpc1*^+/+^, *Mybpc1*^+/-^ and *Mybpc1*^-/-^ newborn mice over first 36 hours of life (n=8-15 per group). I) Similar body development of *Mybpc1*^+/+^, *Mybpc1*^+/-^ and *Mybpc1*^-/-^ at various embryonic stages. **p<0.01 and ***p<0.001. Statistical analyses by one way ANOVA and log rank Mantel-Cox test for (H).

In this study, we generated novel mouse models to systematically investigate the pre- and postnatal cellular and molecular mechanisms through which sMyBP-C regulates skeletal muscle formation and contractile function. Strikingly, both global and muscle-specific conditional knockouts (KO) of *Mybpc1* lead to neonatal lethality within the first day after birth, or in adults, result in compromised muscle formation and impaired muscle function. Our findings demonstrate that sMyBP-C is indispensable for facilitating actomyosin interactions, generating mechanical force and maintaining sarcomere structural organization. Through these pathways, sMyBP-C regulates muscle development and maturation during embryogenesis and facilitates growth and maintenance in postnatal muscle development. In addition to advancing our understanding of the pivotal role played by sMyBP-C, in both the development and function of skeletal muscle, these results hold promise for guiding the formulation of effective therapeutic strategies for related disorders.

## Results

### Early sMyBP-C expression is required for postnatal survival

To investigate the functional and structural roles of sMyBP-C at various stages of muscle development, we examined its expression profile in differentiating C2C12 myoblasts and developing skeletal muscles. sMyBP-C protein was detected within one day after the initiation of differentiation and increased in parallel with myosin heavy chain expression during myotube formation in C2C12 cells. In contrast, fMyBP-C was not detected until approximately one week after initiation of differentiation (Figure 1B and C). A similar pattern of gene expression was observed in developing skeletal muscles, where *Mybpc1* gene expression was detected one week after birth and maintained during muscle growth until eight weeks. By contrast, *Mybpc2* gene expression was low in the early stages but increased significantly at four weeks in EDL and eight weeks in SOL muscles (Figure 1D). These results suggest that early sMyBP-C expression may play a role in the formation and maturation of skeletal muscle structure.

To define the role of sMyBP-C in embryonic musculoskeletal development and survival, we developed the *Mybpc1*gKO mouse model using the CRISPR Cas9 system. This involved switching 859CG to A in exon 8 of the *Mybpc1* gene, which caused a premature stop codon at amino acid 215 (Figure 1E). We confirmed complete deletion of sMyBP-C protein and transcripts in homozygous *Mybpc1*gKO diaphragm muscles (Figure 1F and Suppl. Figure 1). Body weight was slightly but significantly lower in homozygous pups (*Mybpc1g*KO^-/-^) compared to heterozygous (*Mybpc1*gKO^+/-^) and wild-type (*Mybpc1*gKO^+/+^) littermates, and all the *Mybpc1*gKO^-/-^ pups died within 24 h after birth (Figure 1G-H). Developing embryos (E15.5 to E18.5) and newborn mice (P1) revealed significant hypercontractile wrist and severe kyphosis in the *Mybpc1*gKO^-/-^ mice. Deletion of *Mybpc1* also caused severe whole-body tremors and complete immobility after birth (Figure 1I, 2A-C, and Suppl. Video 1).

### Global *Mybpc1* KO causes severe respiratory stress and muscle atrophy

In our investigation of *Mybpc1*gKO^-/-^ mice, we observed abnormal breathing patterns in newborn pups. Plethysmography revealed a significantly lower respiratory rate and irregular breathing patterns in *Mybpc1*gKO^-/-^ mice compared to controls (Figure 2D-F and Suppl. Figure 2B-C). However, tidal volume was not significantly different between the groups, and neuronal development in the diaphragm was preserved in *Mybpc1*gKO^-/-^ (Suppl. Figure 2A, D-F). We also found that contractile functions of the diaphragm were reduced in the *Mybpc1*gKO^-/-^ mice, with significantly lower force generation and calcium sensitivity of skinned muscle, while force redevelopment rate (*k*tr) after slack and re-stretch test was increased in *Mybpc1*gKO^-/-^ compared to controls (Figure 2G-J).

**Figure 2.**
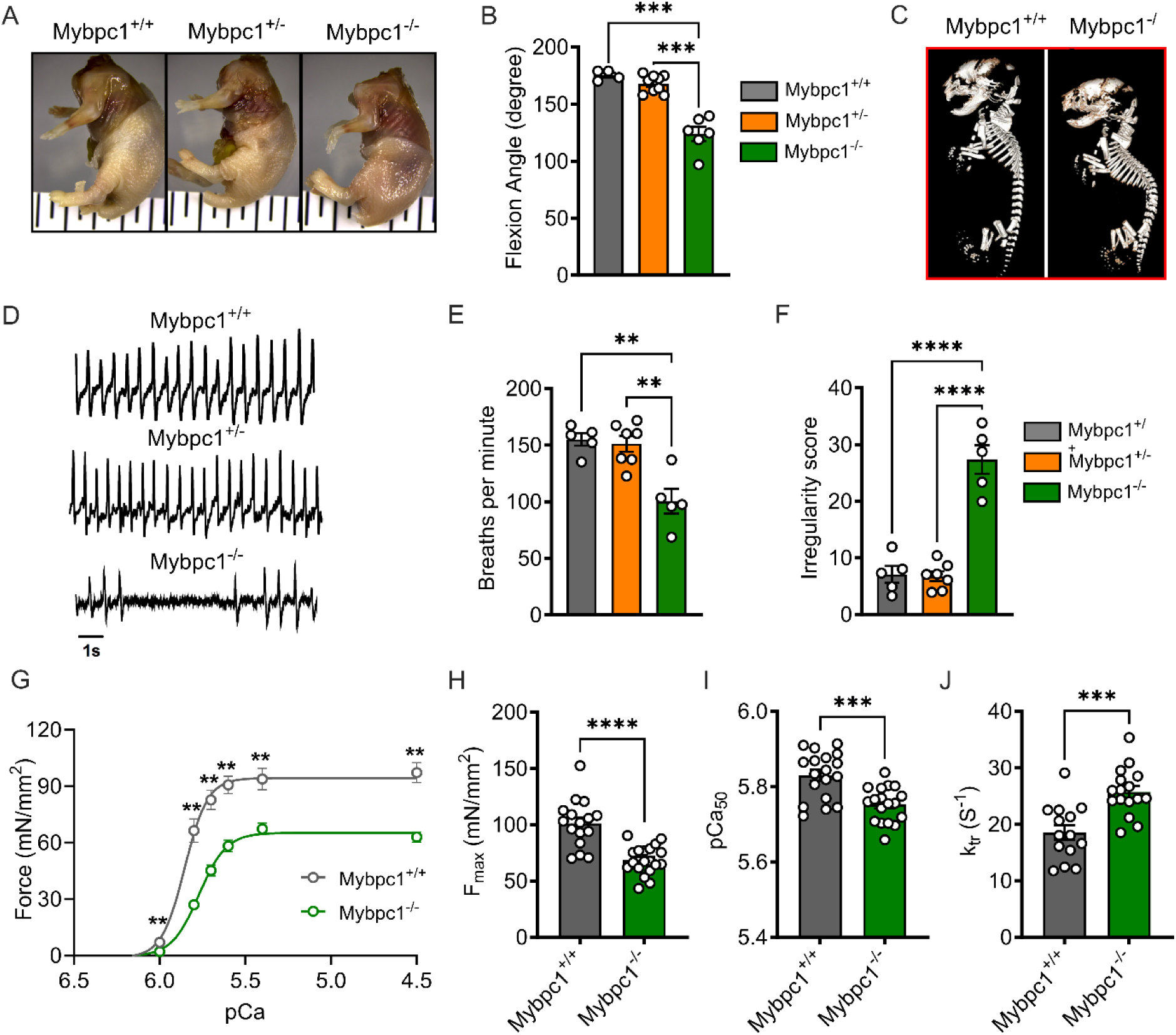
Congenital contractures, respiratory distress and functional deficit after early sMyBP-C deletion. A) Wholemount analysis of wild-type (*Mybpc1*^+/+^), *Mybpc1*gKO^+/-^ (*Mybpc1*^+/-^) and *Mybpc1*gKO^-/-^ (*Mybpc1*^-/-^) newborn mice and B) contracture of the forelimb (n=4-9 per group). C) X-Ray scan of fixed neonatal pups demonstrating kyphosis in *Mybpc1*^-/-^ pups. D) Representative plethysmography traces for *Mybpc1*^+/+^, *Mybpc1*^+/-^ and *Mybpc1*^-/-^ neonatal pups. E) Average number of breaths and F) calculated breath irregularity scores in *Mybpc1*^+/+^, *Mybpc1*^+/-^ and *Mybpc1*^-/-^ pups. G) Skinned diaphragm fiber force-pCa curves with H) maximum force production, I) calcium sensitivity in skinned diaphragm muscle fibers and J) Rate of tension re-development at pCa4.5. **p<0.01, ***p<0.001, p<0.0001. Statistical analyses for (B), (E) and (F) by one way ANOVA and t-test for (G to J).

To elucidate the underlying mechanisms of respiratory stress and functional loss in *Mybpc1*gKO^-/-^ mice, we performed histology and gene expression profiling of the diaphragm muscle. Our analysis revealed a significant reduction in the average size of diaphragm muscle fibers in *Mybpc1*gKO^-/-^ (Figure 3A and B), as well as smaller-sized muscle fibers in hindlimb muscles (Suppl. Figure 2G-I). RNAseq analysis of diaphragm muscle identified a total of 277 differentially expressed genes and 580 related pathways. Gene set enrichment analysis (GSEA) revealed up-regulated pathways associated with muscle atrophy and negative regulation of muscle differentiation while down-regulated pathways included fat metabolism and muscle development (Figure 3C, D and F). Finally, in *Mybpc1*gKO^-/-^ diaphragms, we observed an increase in numerous muscle atrophy-related genes and key genes related to sarcomere structure were dysregulated (Figure 3E and G). In summary, our findings from the *Mybpc1*gKO^-/-^ mouse model demonstrate the essential role of sMyBP-C in both embryonic musculoskeletal development and postnatal survival. The severe skeletal deformity and immature muscle development observed in *Mybpc1*gKO^-/-^ mice leads to perinatal demise, most likely from markedly impaired respiratory function.

**Figure 3.**
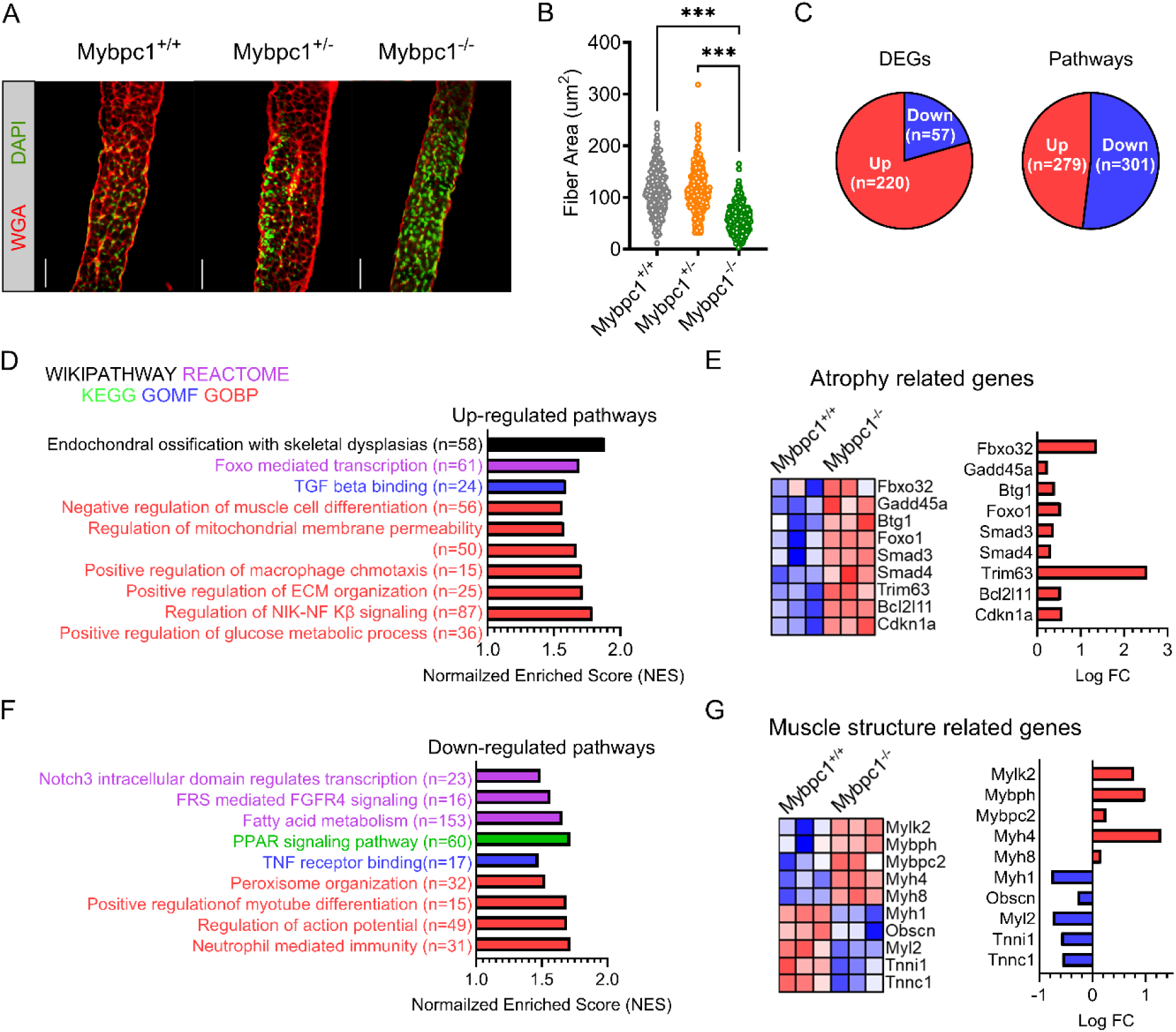
Atrophied muscle and disrupted gene expressions in *Mybpc1* global knockout mice. A) Cross-sections of diaphragms stained with wheat germ agglutinin and DAPI. Scale bar=50um. B) Quantification of myofiber size (from panel A). C) RNAseq analysis of total number of differentially expressed genes and associated pathways in *Mybpc1*^-/-^ diaphragm. Gene Ontology terms related to significantly upregulated (D) and downregulated (F) genes in *Mybpc1*^/-^ diaphragms. Select gene expression related to muscle atrophy (E) and muscle structure (G). ***p<0.001. Statistical analysis by one way ANOVA. Cutoff set for DEGs is logFC>1.5 and p<0.001.

### Impaired muscle growth and functional capacity after *Mybpc1* KO after birth

During postnatal growth, changes occur in skeletal muscle fiber types and key sarcomere proteins, such as myosin heavy chain. To investigate the impact of sMyBP-C on early muscle development and functional capacity after birth, we generated a *Mybpc1* muscle specific conditional KO mouse (*Mybpc1*^fl/fl^MCK^cre^, *Mybpc1*cKO), in which the floxed exon 5 of the *Mybpc1* gene was deleted under the control of the MCK promoter-derived Cre recombinase, which is constitutively active after birth (Figure 4A). At 3 to 4 months of age, sMyBP-C protein was almost completely removed (>99%) in Mybpc1cKO muscles, while fMyBP-C protein levels were upregulated in fast twitch muscles (Figure 4B-C). *Mybpc1*cKO mice had significantly reduced body weight and muscle mass compared to control mice (*Mybpc1*^fl/fl^) (Figure 4D-E). We evaluated *in vivo* skeletal muscle function by measuring running capacity, forelimb grip strength and isometric peak plantarflexor torque generation. All measurements were significantly lower in *Mybpc1*cKO than *Mybpc1*^fl/fl^ mice. During the plantar flexor torque test, the activation rate was preserved, but the relaxation rate was decreased in *Mybpc1*cKO (Figure 4F-J).

**Figure 4.**
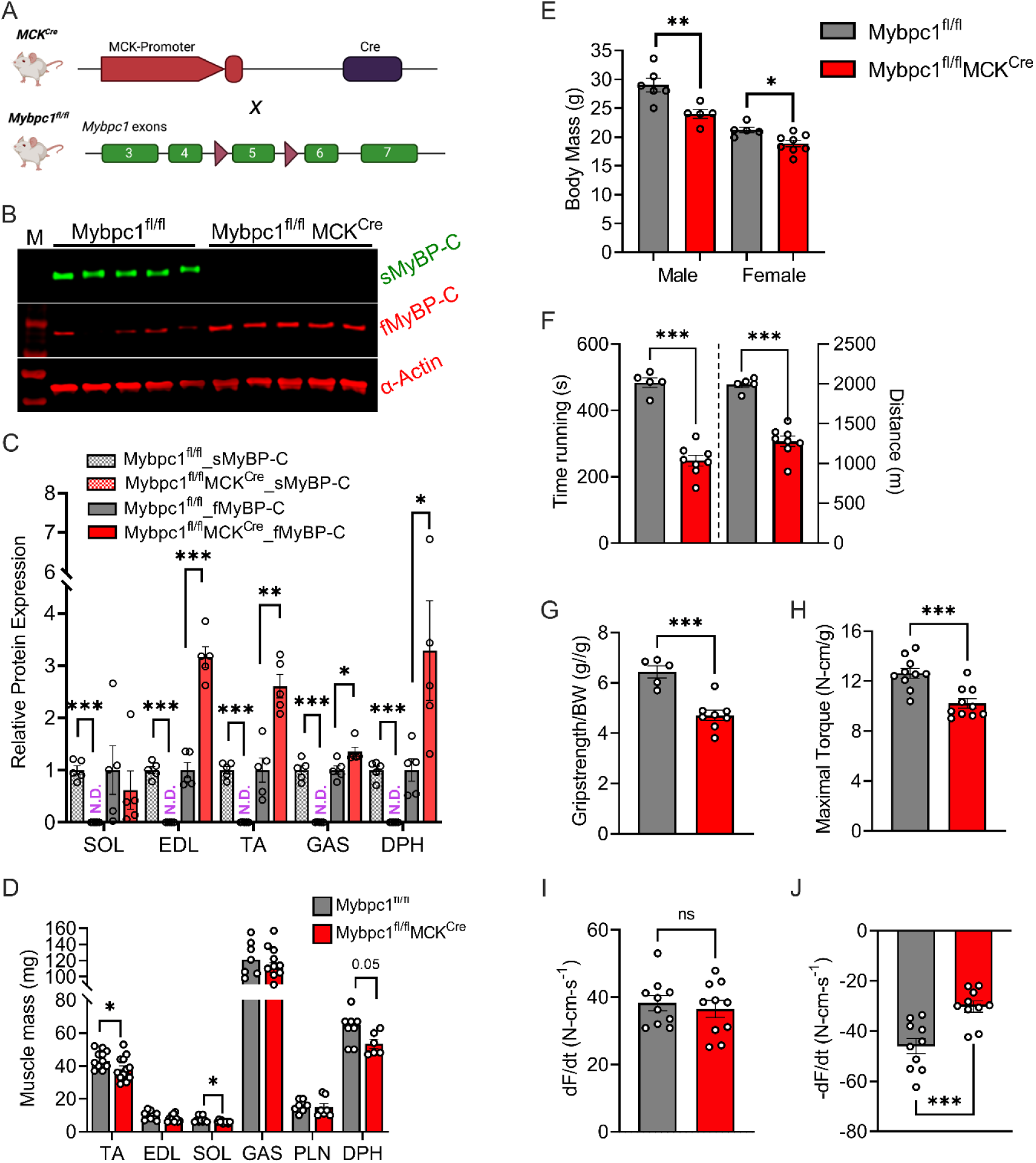
Postnatal deletion of sMyBP-C impairs muscle growth and in vivo muscle function at 3∼4 months old. A) Schematic illustration of skeletal muscle-specific *Mybpc1*^fl/fl^/MCK^Cre^ conditional knockout model. B) Representative Western blot of slow and fast MyBP-C protein expressions in EDL and C) quantification of the two skeletal MyBP-C protein’s expression in slow and fast twitch muscles. D) Muscle mass and E) body weight of 12-week-old mice. *Mybpc1* is required for normal function (n=5-10/group). F) Treadmill running test demonstrating time and distance to exhaustion. G) Grip strength test. In vivo plantarflexion function test showing H) maximal isometric torque production, I) rate of contraction and J) rate of relaxation. *p<0.05, **p<0.01, ***p<0.001. t-test was used for statistical analyses.

To further investigate the function of *Mybpc1*cKO skeletal muscle, we examined soleus muscle *ex vivo* and *in vitro*. Slow twitch type 1 and 2a fibers are dominant in the soleus muscle (*15*) and fMyBP-C does not compensate for the loss of sMyBP-C (Figure 4C), making it an ideal tissue to study the roles of sMyBP-C in skeletal muscle in the absence of compensatory fMyBP-C expression. We found that the isometric peak twitch (Pt) and peak tetanic (Po) force of intact soleus muscle were reduced by 37% and 55%, respectively. Moreover, the specific force (SPo) was significantly decreased, and the half relaxation time (1/2RT) was prolonged in *Mybpc1*cKO, along with slower rates of activation and relaxation, as compared to *Mybpc1*^fl/fl^ (Figure 5A-G). In addition, *Mybpc1*cKO soleus muscle generated less submaximal force at low to medium electrical stimulation and exhibited lower fatigue resistance in response to repeat tetanic muscle contractions compared to *Mybpc1*^fl/fl^. Interestingly, during low frequency tetanic electrical stimulation, we observed that force was unable to accumulate in a manner similar to WT soleus muscle. Instead, after maximal force was generated, an oscillating plateau followed by a loss of force mid-stimulation was observed (Suppl. Figure 4A2). Maximal and submaximal force generation capacities were measured in vitro at the skinned muscle fiber level. The results showed a significant reduction in force generation over a wide range of calcium concentrations in *Mybpc1*cKO fiber, along with a decreased calcium sensitivity and increased *k*tr (Figure 5J-N). Additional functional measurements were performed on fast twitch EDL muscle, revealing a significant reduction in normalized peak tetanic force and specific force, as well as an increase in half relaxation time and a decrease in the rate of relaxation in *Mybpc1*cKO (Suppl. Figure 5).

**Figure 5.**
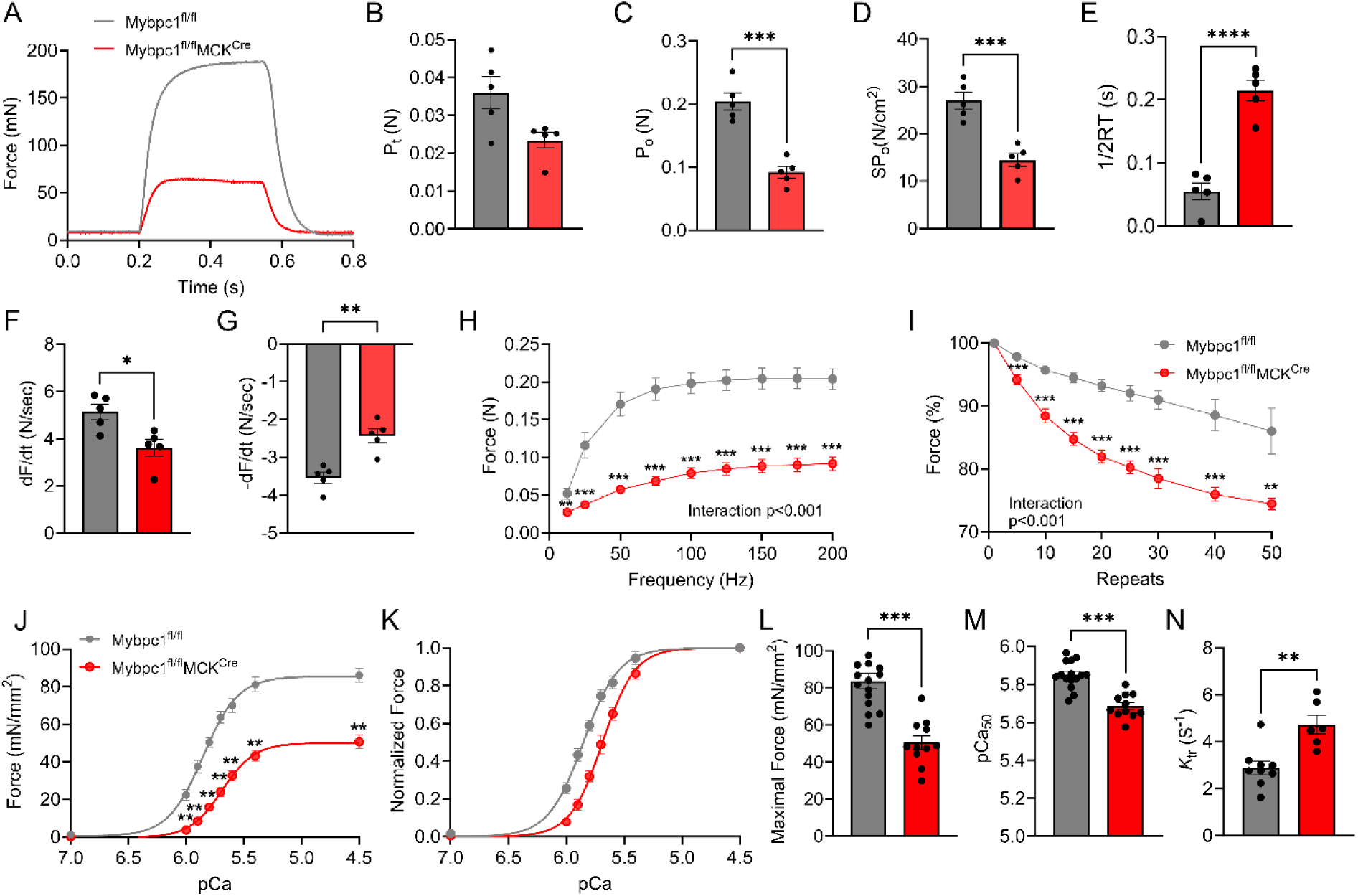
sMyBP-C knockdown after birth reduced contractile functions of slow twitch soleus muscle. A) Representative ex vivo peak isometric tetanic force generation graph. B) Peak twitch force, C) peak tetanic force, and D) specific force generation. E) Half relaxation time, F) rate of activation, and G) rate of relaxation during the peak isometric contraction. H) Force-frequency graph depicted from force generation at electrical frequency at 12.5∼200Hz. I) Fatigue resistance profile after 50 repeated isometric contraction at 150Hz. J) In vitro isometric force generation of skinned SOL fiber from pCa 7.0 to 4.5. K) Normalized force at different calcium concentration. L) Peak isometric force at pCa4.5, M) calcium sensitivity of contraction and N) force re-development rate. *p<0.05, **p<0.01, ***p<0.001, ****p<0.0001 after t-test.

These comprehensive functional analyses, conducted in whole muscle and isolated muscle fibers, consistently demonstrate that sMyBP-C is essential for generating submaximal and maximal force and also for relaxing the muscle after contraction. In addition, sMyBP-C is necessary for maintaining repeated force generation capacity in the slow-twitch soleus muscle.

### Muscle atrophy and fiber type changes in response to conditional *Mybpc1*cKO

To understand the underlying cellular and molecular mechanisms of the functional loss of sMyBP-C in the *Mybpc1*cKO (*Mybpc1*^fl/fl^/MCK^cre^) muscle at 3∼4 months, we investigated the histological adaptations of soleus muscle. We stained cross-sectioned soleus muscle with myosin heavy chain (MHC) antibodies and analyzed fiber size and types. Our results showed a significantly increased number of small-sized fibers but a decreased number of large-sized fibers in *Mybpc1*cKO soleus muscle (Figure 6A-B). Moreover, we observed a near-total elimination of fast-twitch type 2x and 2b fibers, with an increase in the population of type 2a fibers. Additionally, the cross-sectional area (CSA) of all fiber types were significantly smaller, while the number of fibers increased in *Mybpc1*cKO soleus (Figure 6C-E). We also investigated whether muscle atrophy and hyperplasia after *Mybpc1*cKO were due to impaired muscle cell fusion and longitudinal muscle growth. De-membraned soleus muscle fibers were stained with DAPI, followed by measuring the length and number of nuclei in a single soleus fiber. Our results showed no statistical difference in the number of nuclei per millimeter of fiber, indicating that myonuclei fusion during muscle growth after birth was preserved in the KO. However, *Mybpc1*cKO soleus fibers contained significantly more nuclei when normalized by estimated fiber volume with singnifican reduction in fiber diameter (Figure 6F-I). In fast-twitch EDL muscle, the proportion of slow-twitch fibers increased, and the CSA of type 2a fibers decreased in *Mybpc1*cKO. The number of type 2b fibers did not change, but that of other fiber types increased significantly (Suppl. Figure 6). These histology results, which show fiber atrophy and a transition from fast to slow fiber types, provide the cellular mechanisms that underlie the observed functional decline in the *Mybpc1*cKO muscles.

**Figure 6.**
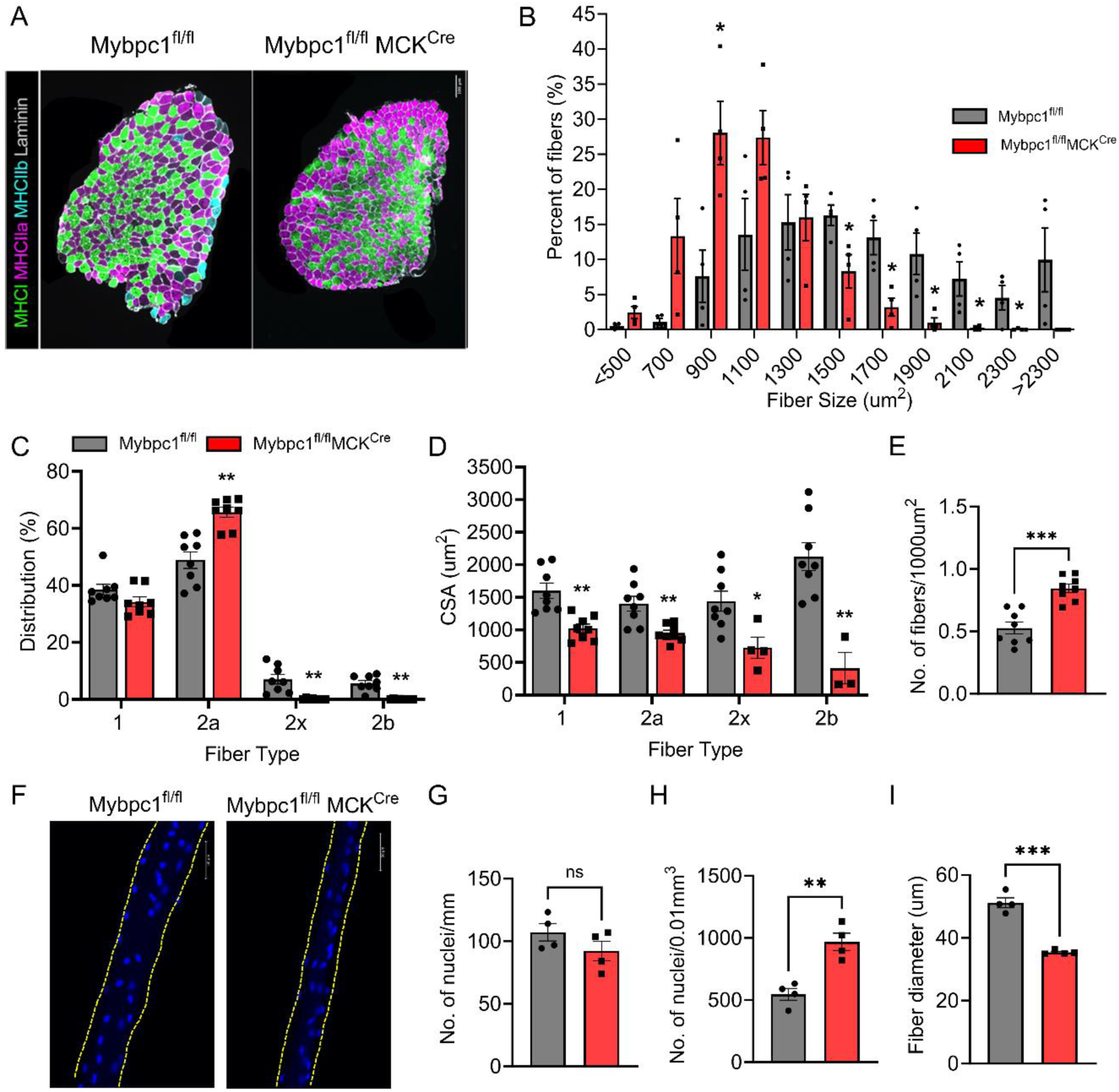
Postnatal sMyBP-C deletion causes muscle atrophy, fiber type switch and disruption of muscle integrity. A) Cross-sectioned soleus samples immunostained with antibodies against myosin heavy chain I (green), myosin heavy chain IIa (magenta), myosin heavy chain IIb (cyan) and laminin (grey). Scale bar=100um. B) Fiber CSA and C) fiber type distribution in soleus muscles. D) Average cross-sectional area of each fiber type and E) numbers of fibers per 1000um^2^. F) Single soleus fiber stained with DAPI. The edge of the fiber is highlighted by dotted lines. Scale bar=50um. Number of myonuclei normalized by G) fiber length and H) volume. I) Averaged fiber dimeter. *p<0.05, **p<0.01, ***p<0.001 after t-test.

RNAseq analysis was conducted to investigate the molecular mechanisms of the functional deficit and muscle atrophy observed after sMyBP-C deletion. We identified 530 differentially expressed genes (DEGs), of which 208 genes were up-regulated, and 322 genes were significantly down-regulated in *Mybpc1*cKO soleus muscle. Gene set enrichment analysis (GSEA) identified 97 differentially regulated pathways (28 up and 69 down). Pathway analysis showed that a group of genes responsible for sarcomere and extracellular matrix (ECM) structure and muscle contraction were down-regulated, while fat metabolism, muscle necrosis, and immune response regulated pathways were activated in *Mybpc1*cKO (Figure 7A-D).

**Figure 7.**
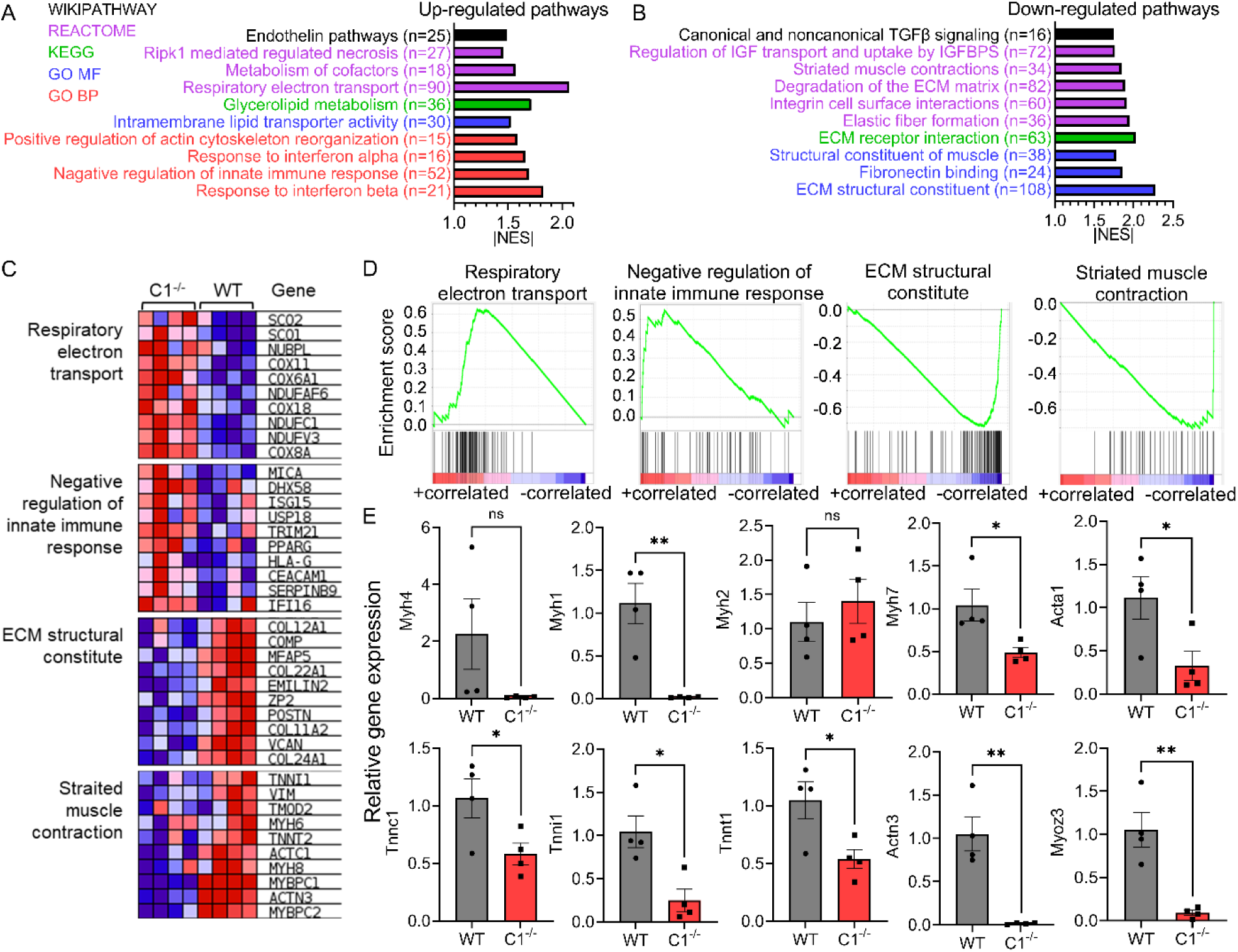
Differential gene expression and molecular pathways in postnatal *Mybpc1*cKO by RNAseq analysis. Ten up (A) or down (B) regulated pathways identified by GSEA in *Mybpc1*^fl/fl^/MCK^Cre^ (C1^-/-^) soleus muscle. C-D) Heatmap and enrichment score graph of key increased or decreased genes of mitochondria respiration, immune response, ECM structure, and muscle contraction. E) qPCR results of key sarcomere genes. DEGs were selected with criteria of logFC>1.5 and FDR<0.05. *p<0.05, **p<0.01. Statistical analyses by t-test for (E).

To confirm the RNAseq results, we examined the expression of key muscle sarcomere, atrophy, and ECM-related genes by RT-qPCR. Consistent with RNAseq results, we found significantly reduced expression of numerous sarcomere genes, such as *Myh1, Mybpc2, Tnnt1, Actc1*, and *Actn3*, as well as troponin complex genes (*Tnnc1, Tnnt1,* and *Tnni1*), Z-disk protein (*Myoz3*), and major skeletal ECM components (*Col1a1* and *Col12a1*) in *Mybpc1*cKO (Figure 7D and Suppl. Figure 7). In particular, gene expression of two key ubiquitin E3 ligases responsible for myofilament proteolysis, Fbxo32 (*Atrogin1*) and Trim63 (*MuRF1*), were significantly reduced in *Mybpc1*cKO, with no changes in *Mstn* and *Foxo1* expressions (Suppl. Figure 7). All the results from histologic and RNAseq analyses presented above demonstrate that sMyBP-C is crucial not only for embryonic muscle development but also for the process of skeletal muscle maturation after birth. In the absence of sMyBP-C, the generation of type 2x and 2b fast twitch fibers is impaired, and the remaining fibers are not able to properly increase in volume or express essential muscle sarcomere and structure components.

### Alterations in sarcomere microstructure in *Mybpc1*cKO soleus muscle

Structural roles of sMyBP-C in soleus muscle were studied by evaluating sarcomere structure and actomyosin interactions using small-angle X-ray diffraction. X-ray diffraction patterns were measured from intact soleus muscle both at rest and during peak isometric tetanic contraction. Interfilament lattice spacing was significantly expanded (∼8%) in resting *Mybpc1*cKO muscle relative to wild type (Figure 8A-D), consistent with the observed reduction in maximum isometric force and increase in *k*tr (*16, 17*). Equatorial intensity ratios (I11/10) were not significantly different between wild type and *Mybpc1*cKO muscle (Figure 8A-D), indicating that there were no differences in radial mass shifts of myosin heads during contraction. Intensity of the meridional reflections including M3, M6, and residual MLL4 under maximum isometric contraction (Figure 8E-G) were all significantly elevated in the absence of sMyBP-C, indicating less recruitment of active myosin heads from the inactive quasi-helically ordered resting configuration in the absence of sMyBP-C during muscle contraction in *Mybpc1*cKO (Figure 8E-G) consistent with reduced isometric force. Axial spacing of the M3 (SM3) and M6 (SM6) meridional reflections was significantly longer in *Mybpc1*cKO than in WT under resting conditions, indicating alterations in thick filament backbone structure, but the expected increase in axial spacing under fully activated conditions was significantly reduced in *Mybpc1*cKO soleus muscle relative to wild type (Figure 8H-K), indicating less strain on the thick filaments.

**Figure 8.**
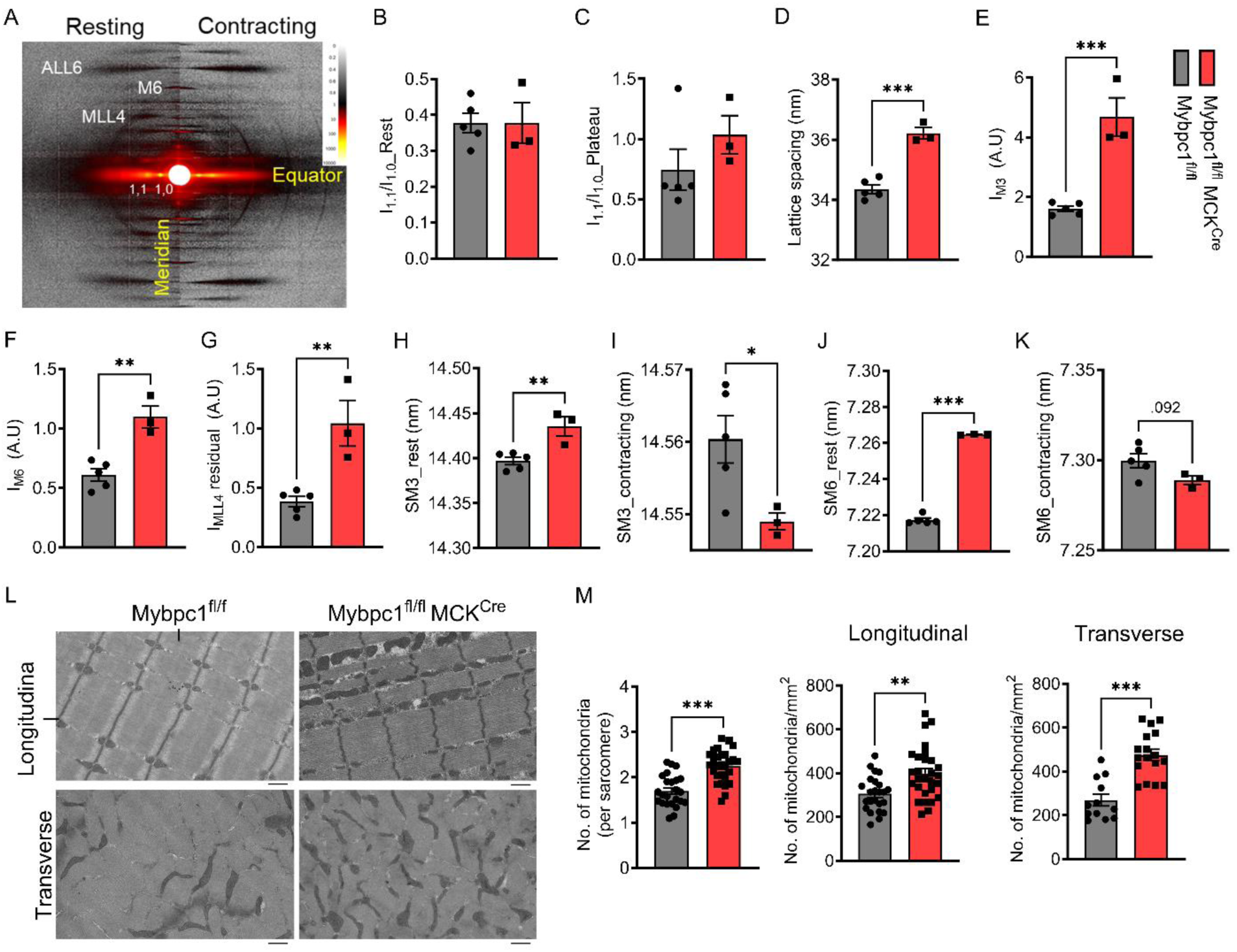
Disrupted sarcomere regulation and structural integrity in soleus muscle after postnatal sMyBP-C deletion. A) Representative small angle X-ray diffraction images in resting and activating conditions. Equator I1.1/1.0 ratio before (B) or during (C) peak isometric tetanic contraction. D) Average lattice spacing between thick and thin filaments. The relative intensity of M3 (E), M6 (F), and residual MLL4 (G) during the peak contraction normalized by its resting values. SM3 and SM6 distances at rest and contracting conditions (H-K). L) Longitudinal and cross-sectioned electron microscopy images of *Mybpc1*^fl/fl^ and *Mybpc1*^fl/fl^/MCK^Cre^ soleus muscle. Scale bar=1um. M) Number of mitochondria were counted and normalized by sarcomere (Left) and area (longitudinal, Middle and transverse, Right). *p<0.05, **p<0.01, ***p<0.001. Averaged value of two groups were compared by t-test.

We further evaluated the structural integrity of sarcomeres lacking sMyBP-C by electron microscopy, which revealed misaligned streaming of the Z-disk in *Mybpc1*cKO (Figure 8L). We also found a significantly increased number of mitochondria in longitudinal and transverse views and within each sarcomere (Figure 8M). Mitochondria were randomly located and frequently found in the space between myofibrils in *Mybpc1*cKO. These results suggest that sMyBP-C is required for maintaining both sarcomere structure and metabolic homeostasis in skeletal muscle.

### Functional deficit and fiber type changes in adult *Mybpc1*cKO

In order to further investigate the roles of sMyBP-C in the function and structure of adult skeletal muscle, we generated a second conditional *Mybpc1*cKO mouse model using a tamoxifen-inducible HSA-merCremer system, achieving over 90% sMyBP-C protein knockdown at three months (Figure 9A-B). Grip strength and running capacity were significantly decreased in the adult *Mybpc1*cKO (*Mybpc1*^fl/fl^/HSA^Cre^) at five months old. In intact soleus muscle, peak isometric tetanic force, specific force and fatigue resistance were also significantly lower in the KO compared to the control (*Mybpc1*^fl/fl^) (Figure 9E-H). We observed similar force reductions in skinned soleus fibers after the conditional KO at maximal and submaximal calcium concentrations, but the calcium sensitivity was preserved in the KO (Suppl. Figure 8B, right). Histological analyses showed significant muscle atrophy (-19% CSA) and fiber type switching (2x and 1 to 2a) in the conditional *Mybpc1*^fl/fl^/HSA^Cre^ soleus muscle. These phenotypes were consistent with those observed in the early postnatal *Mybpc1*cKO (*Mybpc1*^fl/fl^/MCK^Cre^) muscle, indicating that *Mybpc1* not only has essential functional and structural roles in early muscle development but also impacts critical regulatory mechanisms that regulate muscle homeostasis functions into adulthood.

**Figure 9.**
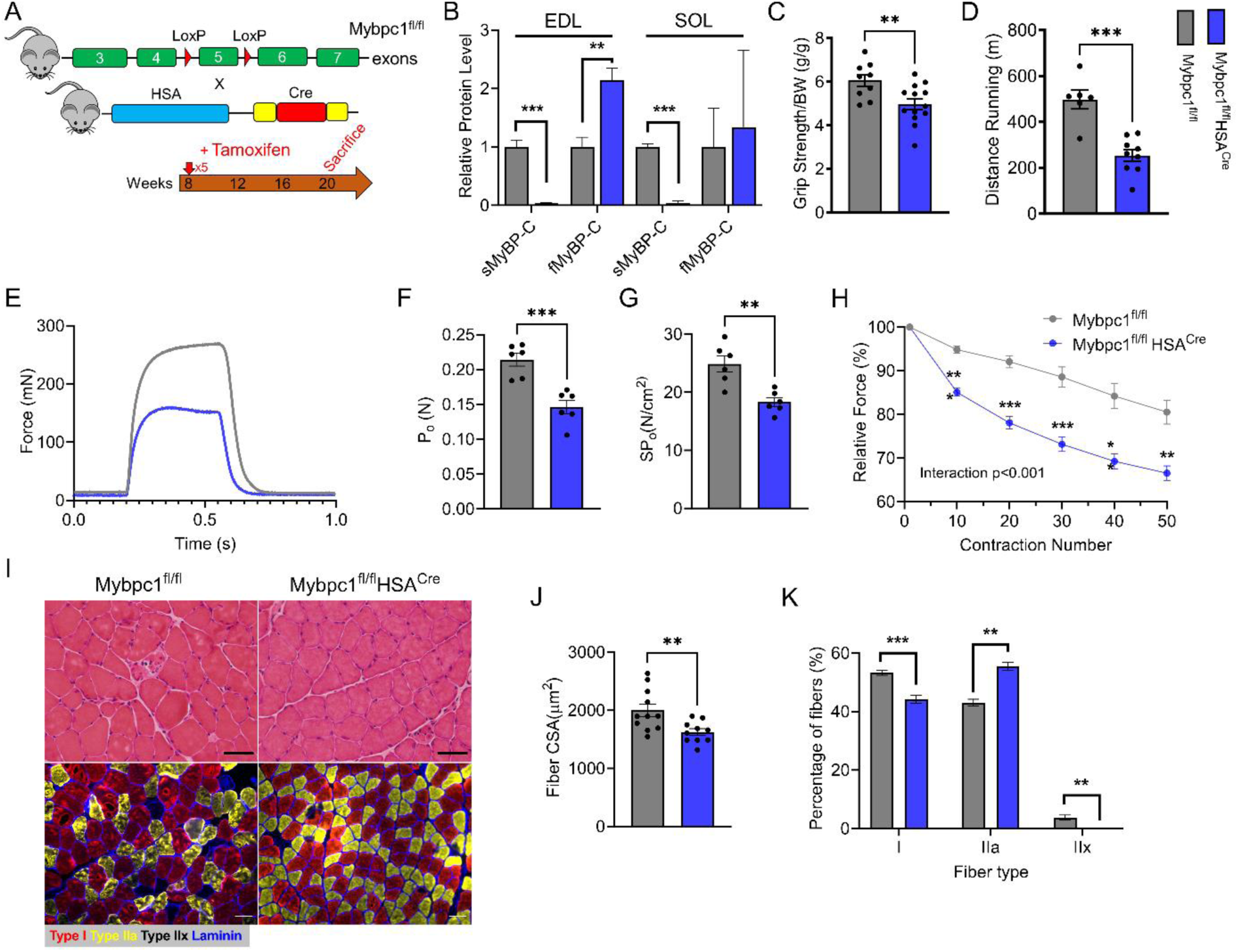
sMyBP-C knockdown at adult stage also compromises muscle function and structure. A) Schematic diagram of skeletal muscle specific adult conditional *Mybpc1*^fl/fl^HSA^Cre^ knockout model. B) Knockdown of sMyBP-C protein expression in EDL and soleus muscles. Grip strength (C) and running capacity (D) were significantly reduced in the KO. Reduced *ex vivo* soleus muscle function; peak isometric tetanic force (E-F), specific force (G), and fatigue resistance (H) in *Mybpc1*^fl/fl^/HSA^Cre^. I) Cross-sectioned soleus samples were stained with H&E (Top) and MHC isoform antibodies (Bottom). Scale bar=50um. Fiber CSA was significantly reduced, and fiber types were switched from type 2X and 1 to type 2A after Mybpc1KO at two months (J-K). **p<0.01, ***p<0.001.t-test was used for statistical analyses.

## Discussion

Mutations in sMyBP-C cause skeletal muscle myopathies, primarily DA (*10, 14, 18–20*). As such, sMyBP-C represents a potentially powerful therapeutic target to understand and reverse contractile deficiencies inherent to DA. Comprising approximately 2% of the myofilament mass, sMyBP-C plays important roles in both contraction and relaxation *in vitro* (21, 22). *In vitro* and *in situ* studies have shown that sMyBP-C is a key player in the regulation of contractility (19, 20, 23, 24). Specifically, sMyBP-C mutations in a Zebrafish model (13) resulted in reduced survival and tail contracture. Investigators have also used *in vivo* electroporation of a CRISPR-mediated knockdown plasmid to assess the role of Mybpc1 in mouse footpad fibers (25, 26). However, these studies only targeted a limited group of muscles, and knockdown was incomplete, making it impossible to discover the true impact of global sMyBP-C ablation (21, 26). In the present study, we systematically unraveled the novel structural, functional and regulatory roles of sMyBP-C at various stages of skeletal muscle development. Our findings reveal that sMyBP-C is expressed early on during myotube formation and skeletal muscle growth. Global Mybpc1 KO experiments have provided compelling evidence that sMyBP-C is indispensable for embryonic musculoskeletal development and postnatal survival. These findings mark a paradigm shift in our understanding of sMyBP-C’s role within skeletal muscle. sMyBP-C has been considered as a sarcomere accessary protein, but we have demonstrated sMyBP-C is vital for skeletal muscle development. In addition, previously unrecognized roles in postnatal muscle function and maturation we found sMyBP-C is involved in congenital muscle diseases, such as arthrogryposis.

Both hetero- and homozygous *Mybpc1*gKO mice were born according to predicted Mendelian expectations. Body weights and sMyBP-C protein levels were normal in heterozygous pups, demonstrating haplosufficient sMyBP-C expression. However, it is interesting that *Mybpc1*^-/-^ showed a severe phenotype manifesting whole-body tremors, as reported previously (*27*). This phenotype expresses a mutant *Mybpc1* in mice (*27*) with complete immobility and reduced body weight at birth, indicating significant impairment of musculoskeletal development during the embryonic stage. The presence of tremor could be explained by increased activation of potassium channels in the muscle fibers (*28*). In our study, the presence of irregular breathing patterns shortly after birth and cyanotic skin discoloration suggest a severe respiratory distress. In contrast, preserved tidal volume in *Mybpc1*gKO suggested that lung functions were not compromised; instead, decreased breathing frequency and apnea might primarily result from diaphragmatic dysfunction. Histologic analysis revealed significantly smaller muscle fiber size in diaphragm and hindlimb muscles, confirming our hypothesis that sMyBP-C, much like myosin heavy chain, is required for the development of embryonic skeletal muscle. RNA-seq analysis revealed differential expression of 277 genes and 580 related signaling pathways in *Mybpc1*gKO diaphragm muscles. Among them, we have highlighted upregulation of muscle atrophy-related genes and pathways, including *Foxo1, Fbxo32* (Atrogin1), and *Trim63* (MuRF-1), along with a decrease in key sarcomere genes essential for embryonic muscle development, such as *Myh1, Myl2, Tnni1,* and *Tnnc1*. These results demonstrate the crucial role of sMyBP-C in both promoting expression of key sarcomere genes and preventing the loss of muscle structure during embryonic muscle development. Interestingly, embryonic Mybpc1 deletion induced a compensatory increase in fast-twitch or adult muscle-specific genes, including *Mybpc2, Myh4*, and *Mylk2*.

To investigate the structural and functional roles of sMyBP-C in postnatal and adult skeletal muscle, we generated two new conditional *Mybpc1*-knockout (cKO) mouse models (*Mybpc1*^fl/fl^/MCK^Cre^ and *Mybpc1*^fl/fl^/HSA^Cre^). In fast-twitch muscles, sMyBP-C and fMyBP-C are expressed together in the A-bands of sarcomeres. Deletion of fMyBP-C has been shown to increase sMyBP-C expression in EDL muscle (*29*). In this study, we found that the inverse is also true. Specifically, sMyBP-C deletion induced compensatory upregulation of fMyBP-C protein expression in fast-twitch muscles, including EDL and TA. However, in soleus muscle, fMyBP-C expression remained at extremely low levels and could not act as a substitute for sMyBP-C deletion, owing to the lack of type 2b fibers. Postnatal knockdown of sMyBP-C, at ∼8 weeks of age, roughly equivalent to a young adult human, resulted in reduced muscle function, body weight, and muscle mass *in vivo*. A comprehensive analysis of isolated soleus muscle function revealed that *Mybpc1*cKO not only reduced force generation capacity and contractility but also specific force and fatigue resistance, collectively suggesting a decline in muscle quality and integrity. Furthermore, a significant decrease in tetanic force accumulation after each electrical stimulation was observed, and the generated peak force was not sustained during contraction, indicating reduced cross-bridge cooperation and affinity, respectively.

Mechanical tension and force generation play critical roles in skeletal muscle development, particularly at embryonic stages during which appropriate musculoskeletal development requires body movement and muscle growth (*30–33*). Deletion of embryonic sMyBP-C may disrupt actomyosin interactions and impair force generation which would inhibit the expression of key thin and thick filaments, but activate muscle atrophy signaling pathways. To rule out the possibility of neuronal defect, we confirmed proper neuromuscular junctions and acetylcholine receptor development. Therefore, it was concluded that impaired diaphragmatic development and function can cause severe respiratory burden, ultimately leading to the death of newborns, as demonstrated in several preclinical and clinical studies (*34–36*). We also found altered patterns of muscle relaxation in soleus muscle. After isometric tetanic contraction in WT, muscle relaxation was initiated by a slow linear phase followed by fast exponential relaxation. However, in *Mybpc1*cKO muscle, force decreased immediately after cessation of electrical stimulation, and was accompanied by a nearly complete absence of slow relaxation, indicating fast detachment of cross-bridges (*37, 38*). Moon et al. (*39*) reported that MyBP-C could reduce the inhibitory effect of tropomyosin via partial dislocation of the protein, thereby facilitating actomyosin binding and increasing calcium sensitivity. Therefore, conditional knockout of sMyBP-C may disrupt normal regulation of thin filaments, leading to unstable and less cooperative actomyosin interactions and cross-bridge cycles. These mechanical perturbations contribute significantly to delayed maturation and growth in the soleus muscle. Our RNA-seq and qPCR data further confirmed immature skeletal muscle development accompanied by significantly lower expression of sarcomere and extracellular matrix genes, such as *Actc1, Tnnc1, Tnnt1, Tnni1, Col1a1*, and *Col12a1*. An unexpected reduction in the expression of two key ubiquitin E3 ligase genes, *Fbxo32* and *Trim63*, may represent a compensatory response to impaired muscle growth, similar to the observed increase in fiber number in soleus muscle. In fast-twitch EDL, however, upregulation of fMyBP-C could partially compensate for functional loss after deletion of sMyBP-C. Peak twitch force was preserved, and loss of peak tetanic and specific force was milder in EDL compared to soleus muscle.

Our X-ray diffraction results revealed significant changes in sarcomere structure in *Mybpc1*cKO muscle that could explain reduced force output. The larger lattice spacing in *Mybpc1*cKO muscle strongly suggests that sMyBP-C plays a crucial role in anchoring actin and myosin and maintaining optimum distance between thick and thin filaments. The longer thick filament backbone periodicities in *Mybpc1*cKO muscle under resting conditions indicate alterations in thick filament backbone structure in the absence of sMyBP-C. Recent cryo-EM studies also support the role of MyBP-C in stabilizing thick filament backbone structure (*40, 41*). The reduced extension of *Mybpc1*cKO thick filament backbones relative to WT under contracting conditions could result simply from the reduced isometric force or if the changes in backbone structure made the thick filaments stiffer. The observed reduced isometric force can be explained by the smaller fraction of active force-producing heads, as indicated by increased residual MLL4 intensity. Our X-ray diffraction analysis suggests that the absence of sMyBP-C results in expansion of the myofilament lattice and changes in thick filament backbone structure. Together, these effects may cause impaired recruitment of active myosin heads, providing an explanation for reduced maximum isometric force during contraction.

While gross sarcomere structure was preserved in *Mybpc1*cKO, muscle ultra-structure examined with electron microscopy revealed distortion of Z-disk alignment to a wavy shape. Underdeveloped sarcomere structure may be the cause of Z-disk deformation, as evidenced by the decreased expression of sarcomeric genes, such as Myoz3 and Actn3. In addition, the higher number of mitochondria observed in *Mybpc1*cKO is consistent with the increase in respiratory electron transport pathway via RNA-seq analysis. These findings suggest that disruption of energy metabolism and balance may occur in *Mybpc1*cKO mice. Similar Z-disk disarray and increased mitochondria number have been reported in the knockout of cMyBP-C (*42*). Compared to developmental KO of *Mybpc1*cKO (Mybpc1^fl/fl^/MCK^Cre^, adult muscle knockout of *Mybpc1* (*Mybpc1*^fl/fl^/HSA^Cre^) exhibited similar functional and histological phenotypes, including reduced *in vivo* and *ex vivo* force generation, muscle atrophy, and fiber type switch. These results indicate that sMyBP-C is essential in regulating actomyosin interaction and force generation during early muscle development and adult muscle homeostasis.

In summary, our comprehensive and mechanistic assessment demonstrates that sMyBP-C is a vital sarcomere protein necessary for proper muscle growth and function during prenatal, perinatal, and postnatal development, as well as adult stages. sMyBP-C is not only critical for embryonic musculoskeletal development, essential for survival after birth, but also for postnatal muscle growth and homeostasis. The findings of this study make a significant contribution to our understanding of the molecular mechanisms underlying muscle contraction and relaxation and contribute to identifying the genetic disorders that cause muscle diseases associated with mutations of the *MYBPC1* gene.

## Materials and Methods

An expanded Materials and Methods section can be found in the online data supplement available.

### Mouse models

Various mouse models were used in the present study to determine the role of sMyBP-C during embryonic, early post-natal and adult stages in skeletal muscle formation and function. These mouse models include global *Mybpc1* knockout (*Mybpc1*gKO), post-natal conditional *Mybpc1* knockout (*Mybpc1*^fl/fl^/MCK^Cre^) and *Mybpc1* adult conditional knockout (*Mybpc1*^fl/fl^/HSA^merCremer^). All mice were anesthetized by 1.5-2.0% isoflurane inhalation and euthanized by cervical dislocation prior to tissue collection. All animal procedures were performed in accordance with protocols approved by the Institutional Animal Care and Use Committee at the University of Cincinnati.

### Cellular, Molecular, Structural and Functional Analyses

Molecular analyses, including protein and RNA analyses, functional analyses such as plethysmography, treadmill running and grip strength tests, in vivo hindlimb muscle function, ex vivo intact muscle function test, and in vitro skinned fiber test and imaging analyses such as immunohistochemistry, whole-body X-ray scanning, muscle X-ray diffraction, transmission electron microscopy were systematically performed using the diaphragm, soleus, and EDL muscles from the various *Mybpc1* KO mouse models.

### In vitro Cell Culture Studies

C2C12 myoblasts and immortalized myoblasts from wild-type and *Mybpc1*gKO mice were differentiated into myotubes to determine the critical role of sMyBP-C in sarcomere and myotube formation.

### Statistical Analysis

All data is presented as mean ± SE. For group comparisons, we used Student’s t-test or one way ANOVA with Tukey’s post-hoc test using GraphPad Prism 7.04 software. Survival curves of new-born pups were analyzed by Mantel-Cox test. Statistical significance was defined as a P-value less than 0.05.

## Acknowledgments

This work was supported by funds from the National Institute of Arthritis and Musculoskeletal and Skin Diseases grant mechanism (to S.S; R01AR078001). In addition, other funding support was utilized from the American Heart Association Career Development Award (to T.S; 23CDA1046498), National Institutes of Health grants (to S.S; R01AR079435, R01AR079477, R01HL105826, and R01HL130356), American Heart Association Transformation Awards (to S.S; 19TPA34830084 and 945748), National Institutes of Health grants (to J.R.P; HL160966 and R21AR077802), and American Heart Association Predoctoral Fellowship (to M.L.V; 2021AHAPRE216237). This research also used resources at the Advanced Photon Source, a U.S. Department of Energy (DOE) Office of Science User Facility operated for the DOE Office of Science by Argonne National Laboratory under Contract No. DE-AC02-06CH11357. This project is supported by grant P30 GM138395 from the National Institute of General Medical Sciences of the National Institutes of Health. The content is solely the authors’ responsibility and does not necessarily reflect the official views of the National Institute of General Medical Sciences or the National Institutes of Health.

## Supplementary Materials

**Supplemental Figure 1.**
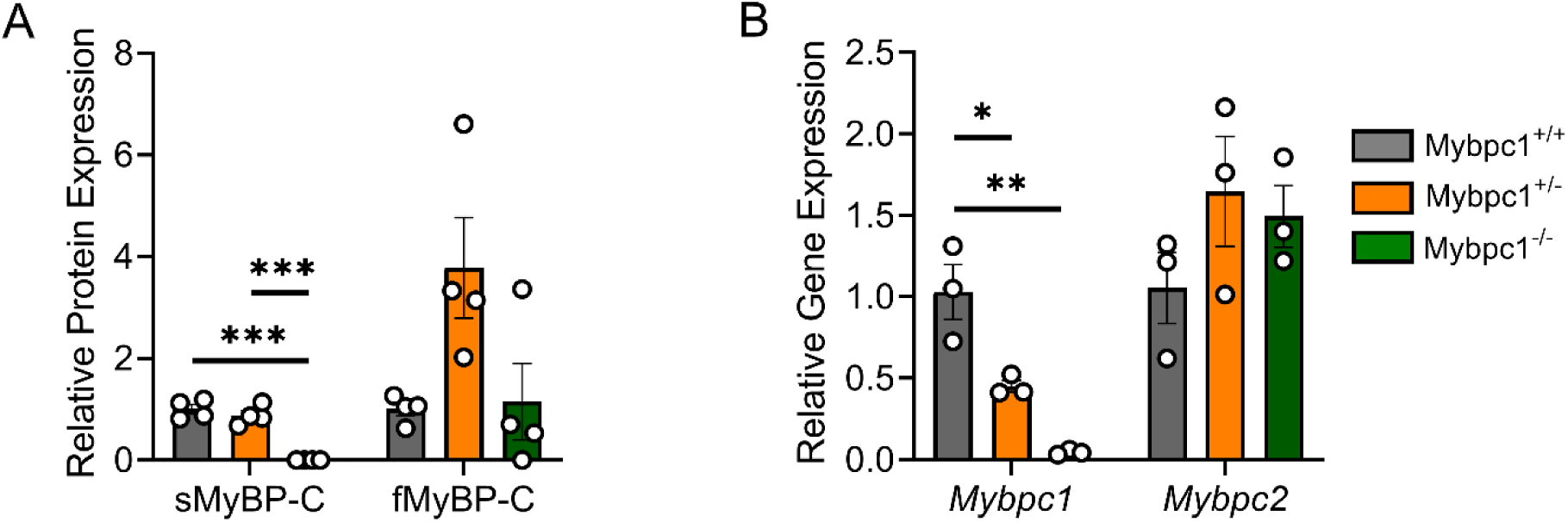
Slow and fast MyBP-C protein and gene expressions in global *Mybpc1*KO muscle. A) Quantification of sMyBP-C and fMyBP-C protein expression in diaphragm muscles by Western blot (n=4 per group). B) qPCR analysis of *Mybpc1* and *Mybpc2* transcript levels in hindlimb samples (n=3 per group). *p<0.05, **p<0.01, ***p<0.001 by one way ANOVA.

**Supplemental Figure 2.**
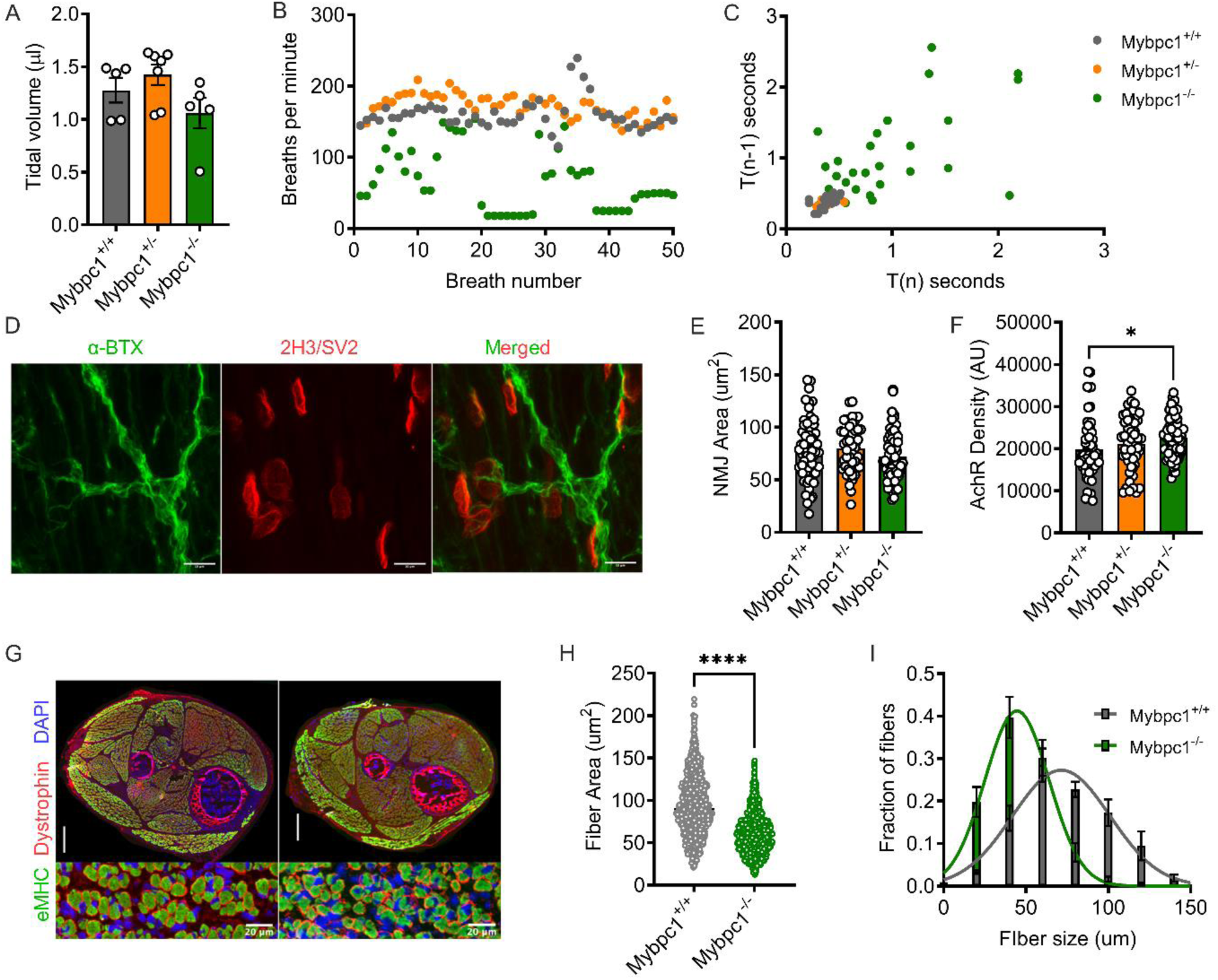
Respiratory function measurement and histology of neuromuscular junction and hindlimb of *Mybpc1*gKO. A) Measurement of tidal volume (n=5∼7 pups). Representative graph depicting breath number (B) and variability (C). D) Immunostaining image of neuromuscular junction (NMJ) with α-bungarotoxin and 2H3/SV2 antibodies. Scale bar=10um. Quantification of NJM area (E) and density of acetylcholine receptor (F). G) Cross-sectioned hindlimbs of *Mybpc1*^+/+^ (left) and *Mybpc1*^-/-^ (right) were immunostained with eMHC, dystrophin and DAPI. Scale bar=300um (top) and 20um (bottom). Averaged CSA (H) and size distribution (I) of hindlimb muscle fibers. **p<0.01, p<0.0001. Statistical analyses for (A), (E) and (F) by one way ANOVA and t-test for (H).

**Supplemental Figure 3.**
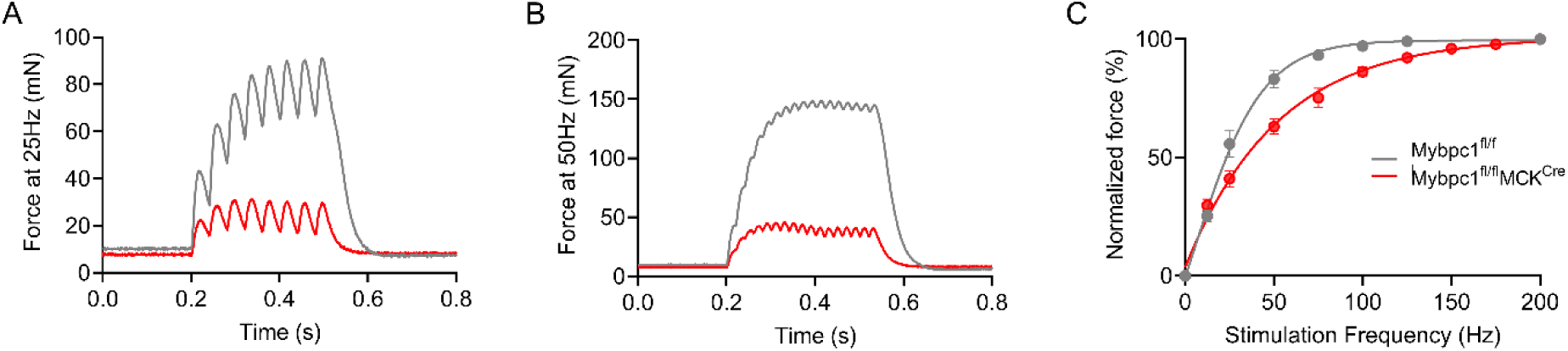
Reduced force generation and cooperativity in Mybpc1cKO soleus muscle. Representative isometric tetanic force generation graph at 25 and 50 Hz electrical stimulation (A and B). C) Down shifted relative isometric tetanic force-frequency graph in *Mybpc1*cKO.

**Supplemental Figure 4.**
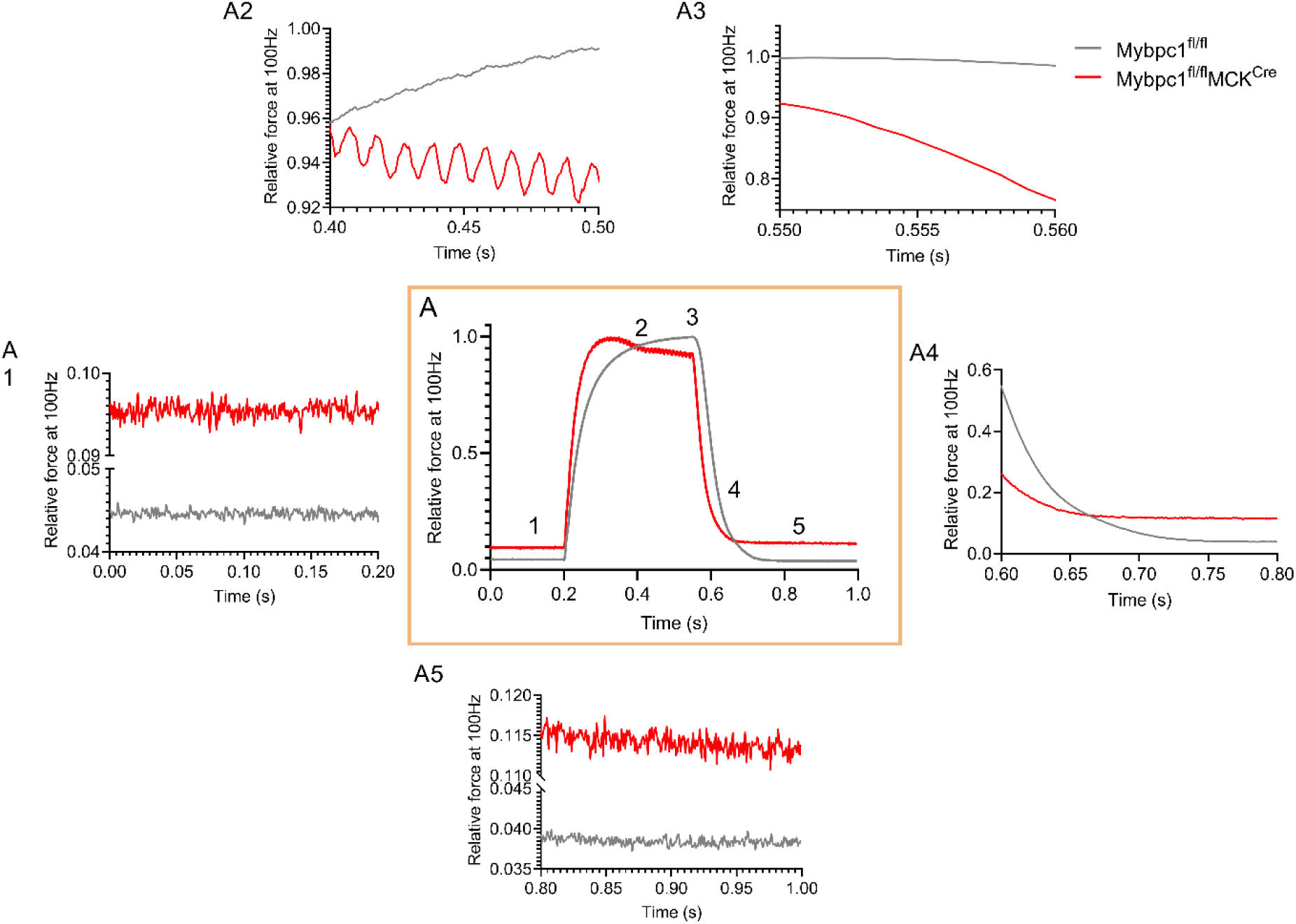
Impaired contraction and relaxation during isometric tetanic contraction. A) Averaged relative isometric tetanic force graph at 100 Hz (center, n=5 in each). Distinct patterns of the force generation graph are highlighted during different periods of contractions (resting, contracting, relaxing1, relaxing2, and resting, A1∼A5, respectively).

**Supplemental Figure 5.**
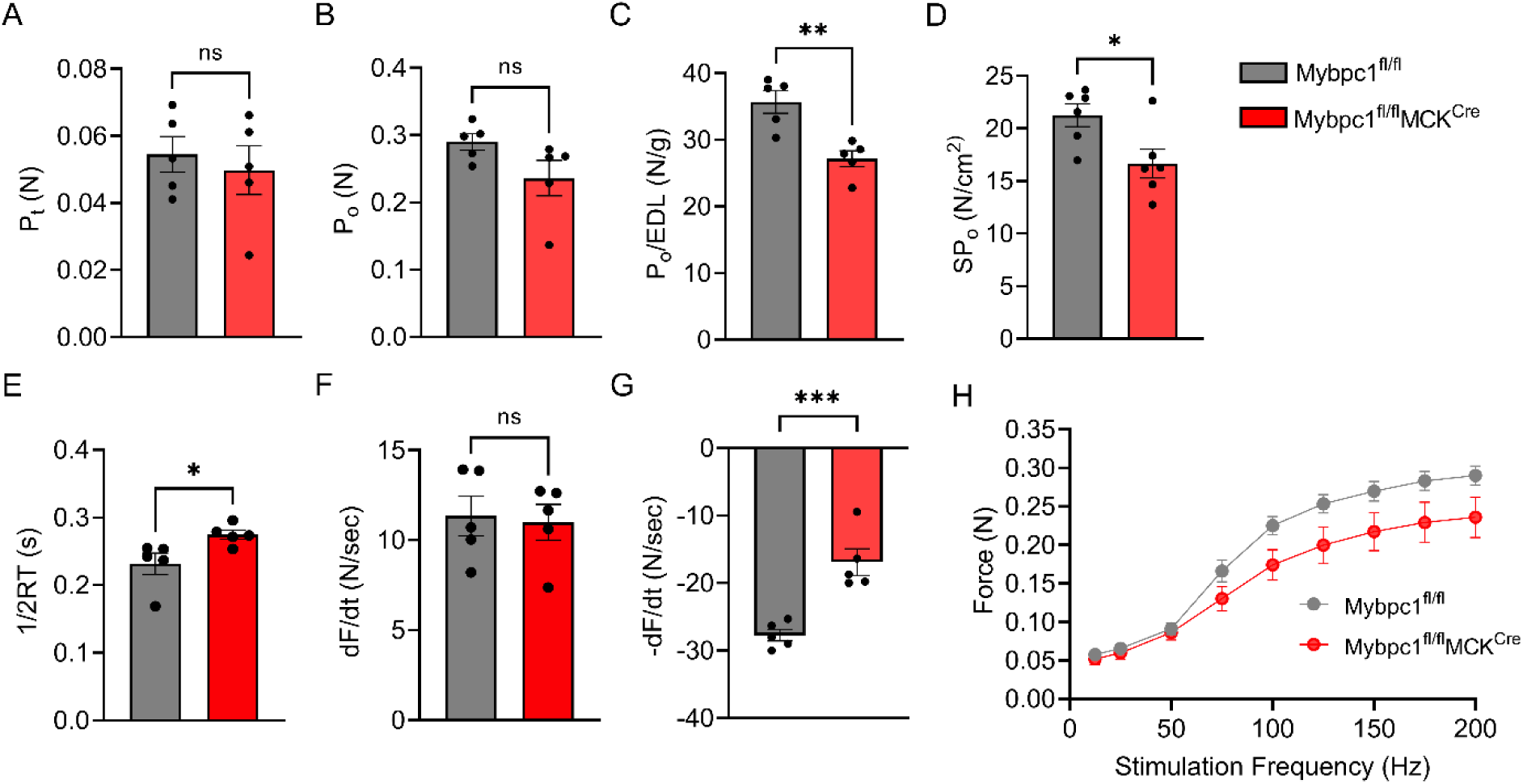
Decreased contractile functions of Mybpc1cKO EDL muscle. A) Peak twitch force (A), peak tetanic force (B), relative peak tetanic force (C) and specific force (D) generation of EDL muscle. Measurements of half relaxation time (E), rate of activation (F), and relaxation (G) during the peak tetanic contraction. H) Absolute force-frequency relation during the isometric tetanic contractions at 12.5∼200Hz electrical stimulation (n=5 in each group). *p<0.05, **p<0.01, ***p<0.001 after t-test.

**Supplemental Figure 6.**
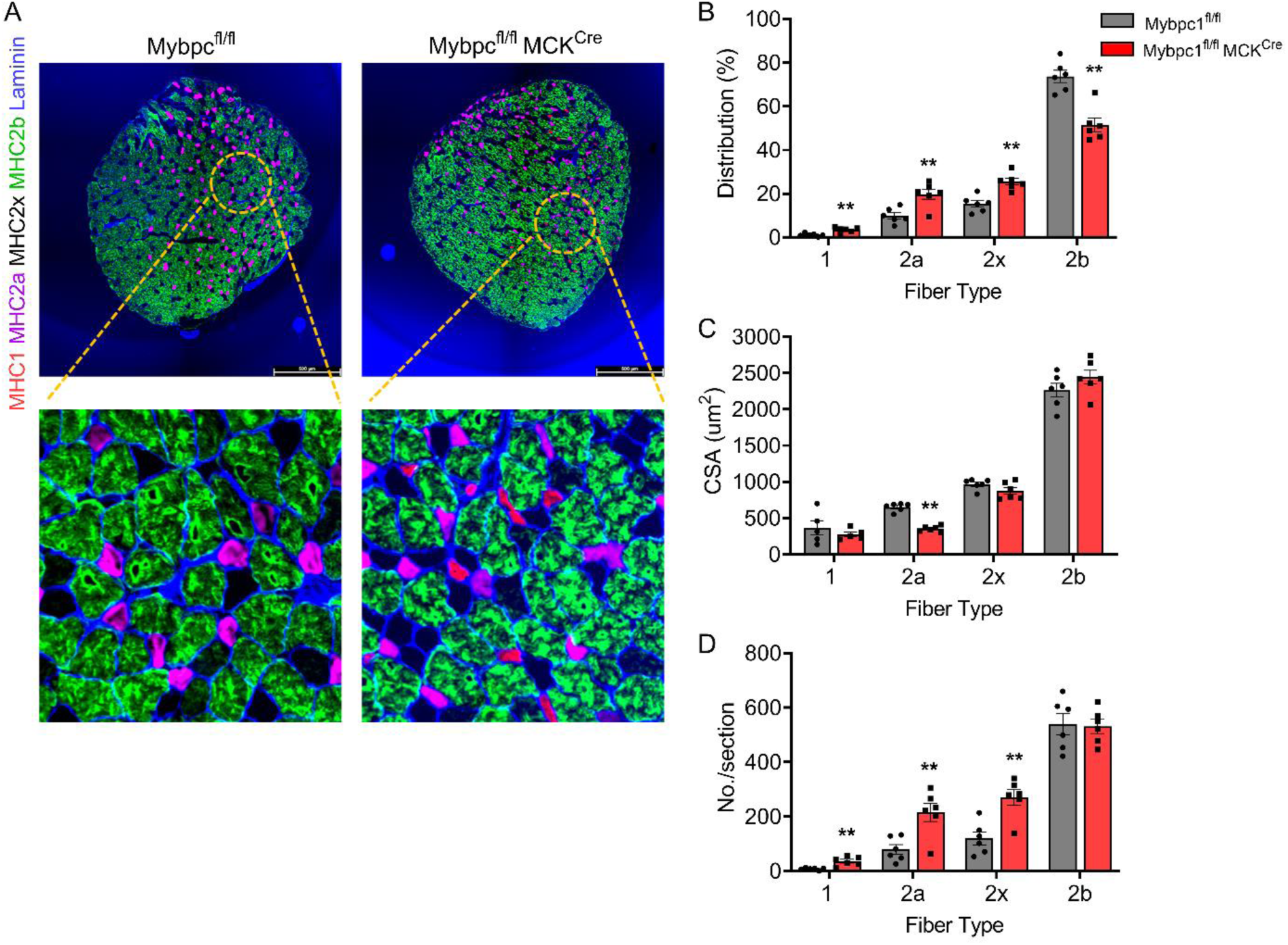
Fiber type switch and atrophy of EDL muscle fiber in *Mybpc1*cKO. A) Representative cross-sectioned EDL muscle immunostained with myosin heavy chain I (green), myosin heavy chain IIa (magenta), myosin heavy chain IIb (cyan) and laminin (blue) antibodies. Scale bar=500um. Fiber type distribution (B), averaged CSA of each fiber type (C) and numbers of each fiber type (D) were quantified from six slides per group, with three mice in each group. **p<0.01 after t-test.

**Supplemental Figure 7.**
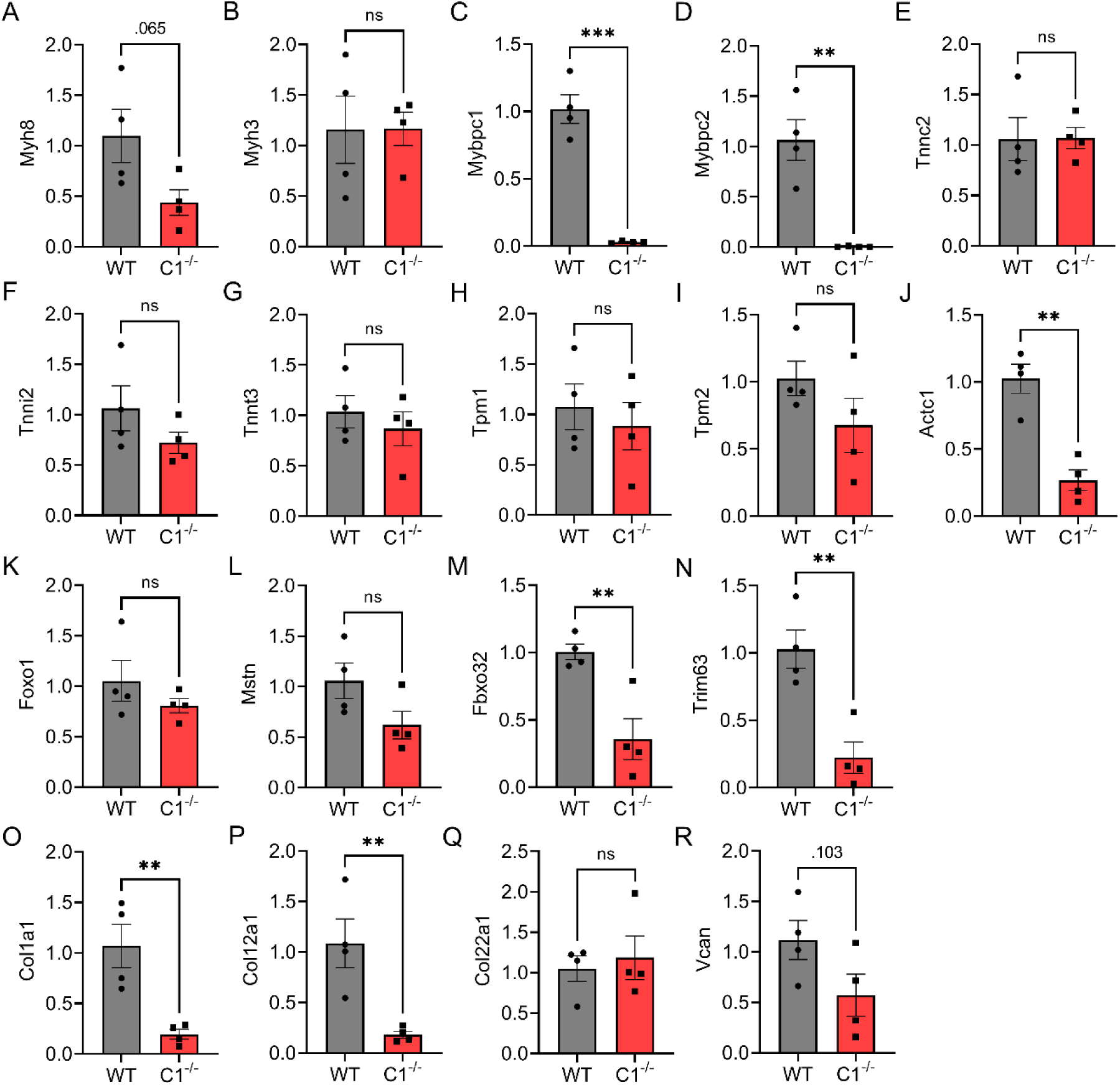
Quantification of gene expression related with sarcomere structure, muscle atrophy and ECM in soleus muscle. Relative mRNA expressions of sarcomere thick and thin filament (A-J), key muscle atrophy (K-N) and skeletal ECM (O-R) were measured by qPCR and compared between *Mybpc1*^fl/fl^ and *Mybpc1*^fl/fl^/MCK^Cre^ (C1^-/-^). Gene expression was normalized by GAPDH (n=4 in each group). **p<0.01, ***p<0.001 after t-test.

**Supplemental Figure 8.**
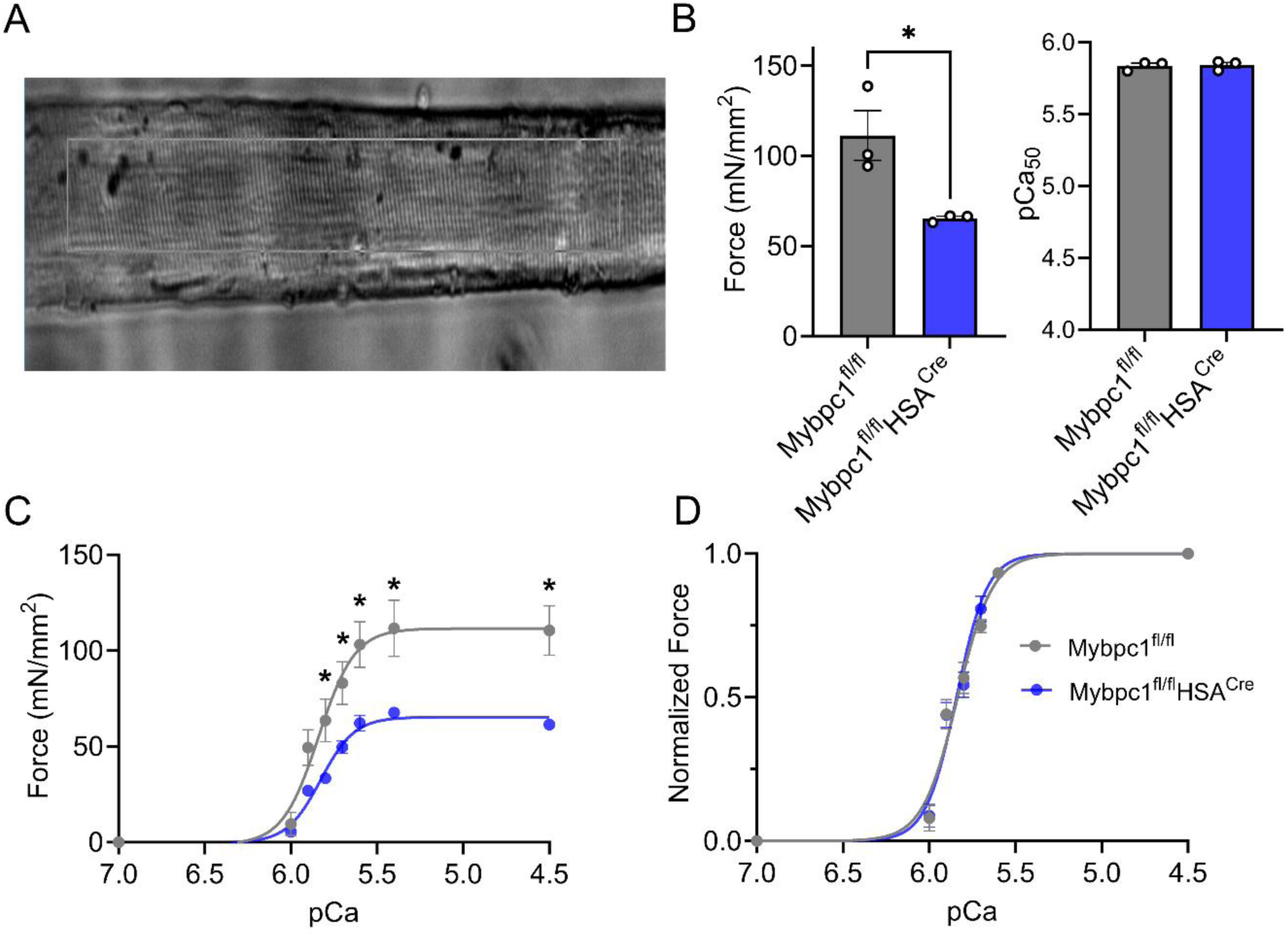
Reduced force generation capacity of skinned soleus fiber in adult *Mybpc1*cKO (*Mybpc1*^fl/fl^ HSA^Cre^) mice. A) Representative picture of skinned single soleus muscle fiber. B) Peak isometric force generation and calcium sensitivity. C) Absolute in vitro isometric force-pCa graph of skinned soleus fiber ranging from pCa 7.0 to 4.5. D) Normalized force generation of the skinned fiber. N=3 in each group. *p<0.05 after t-test.

