## Supplemental Materials for "Unlocking the Role of sMyBP-C: A Key Player in Skeletal Muscle Development and Growth"

### Materials and Methods

C2C12 cell culture

C2C12 myoblasts were cultured in proliferation media (DMEM, 10% fetal bovine serum, and 1% Pen/Strep) until reaching 80~90% of confluence. Then, the culture media was switched to differentiation media (DMEM, 2% horse serum, and 1% Pen/Strep) and replenished every other day. Samples were imaged and collected at 0, 1, 2, 4, 5, 6, 7, 8, 10, 12, and 14 days after differentiation to profile the expressions of myosin heavy chain and skeletal MyBP-C proteins.

Mouse models

*Mybpc1 global knockout (Mybpc1gKO)*

The CRISPR Cas9 system was used to generate a global Mybpc1 KO mouse model. In exon 8 of the Mybpc1 gene, the nucleotides at position 859 were switched from CG to A. This change results in premature termination at amino acid 215, resulting in nonsense mediated decay of the *Mybpc1* transcript.

*Mybpc1 conditional knockout (Mybpc1$^{fl/fl}$ MCK$^{Cre}$)*

To achieve the skeletal muscle-specific Mybpc1 KO immediately after birth, we first created a Mybpc1$^{fl/fl}$ mouse line in which LoxP sites were inserted intron 5 and 6. Muscle creatine kinase - Cre (MCK$^{Cre}$) transgenic mouse (*1*) was crossed with Mybpc1$^{fl/fl}$ mouse containing the floxed exon 5 of Mybpc1 gene, allowing for efficient Mybpc1 deletion by MCK promoter-mediated Cre recombinase after birth.

*Mybpc1 adult conditional knockout (Mybpc1$^{fl/fl}$ HSA$^{merCremer}$)*

We generated conditional adult skeletal muscle specific Mybpc1KO by crossing Mybpc1$^{fl/fl}$ mouse line with HSA$^{merCremer}$ mouse (*2*). Human sarcomere actin (HSA) promoter is conditionally active in skeletal muscle upon tamoxifen treatment. Thus, tamoxifen (80 mg/kg BW) was delivered for five consecutive days via I.P injection at 8 weeks old.

All mice were anesthetized 1.5-2.0% isoflurane inhalation and euthanized by cervical dislocation prior to tissue sample collection. All animal procedures were performed in accordance with protocols approved by the Institutional Animal Care and Use Committee at the University of Cincinnati.

Protein Analyses

C2C12 cells and skeletal muscle samples were homogenized in protein solubilization buffer (PSB, 1632145, Bio-Rad) and the protein concentration was determined by Bradford assay (23238, Thermo Scientific). Subsequently, 10~20 µg of samples was loaded onto pre-cast gradient protean gels (Bio-Rad) and electrophoresed at 100V for 90 minutes. The proteins were then transferred onto nitrocellulose membranes and blocked in 5% non-fat dry milk in Tris-buffered saline containing 0.1% Tween-20 (TBST) for 1 hour. The membranes were incubated with primary antibodies against myosin heavy chain (pan MYH, MF20; DSHB) and MyBP-Cs (sMyBP-C, WH0004604M1 and fMyBP-C, SAB2108180; Sigma-Aldrich) overnight at 4°C. After incubating with Infrared Fluorescent Dyes (IRDye, Licor) or HRP conjugated secondary antibodies for 1 hour at room temperature, the protein bands were visualized and qunatified using the Odyssey CLx (Li-Cor) or Bio-Rad ChemiDoc system. GAPDH or α-actin was used as an internal loading control.

Quantitative RT-PCR

Total RNA was isolated from the snap-frozen skeletal muscle samples using the RNeasy Mini Kit (74104, Qiagen) coupled with an in-column DNase1 treatment (79254, Qiagen). Reverse transcription was performed using the iScript cDNA synthesis kit with a mix of random and oligo-dT primers (1708890, Bio-Rrad Laboratories). mRNA expression was quantified using iTaq Universal SYBR Green Supermix (175120, BioRad) and gene-specific primers. The qPCR was performed on a CFX96 real-time system (Bio-Rad). All reactions were performed in triplicate. Relative gene expression was calculated using the $\Delta\Delta C_t$ method normalized to the GAPDH levels.

Immunohistochemistry

Isolated diaphragm, soleus, and EDL muscles were embedded in O.C.T. Samples were cross sectioned at 10 μm thickness and stained with H&E or specific antibody for immunohistology as previously described (*3*). In brief, slides were fixed with cold acetone for 5 min and permeabilized in 0.5% Triton X-100 in PBS for 20 min. After blocking for 30 min at RT with blocking solution (BlockAid, B10710, Thermo Fisher Scientific), primary antibodies against myosin heavy chain isoforms (MHC type I,BA-D5; MHC type IIA, SC-71; MHC type IIb, BF-F3; Developmental Studies Hybridoma Bank, University of Iowa), embryonic myosin heavy chain (eMHC, HPA021808, Sigma-Aldrich), α-Bungarotoxin (B13423, Thermo Fisher), 2H3/SV2 (SV2, DSHB), WGA (W11261, Thermo Fisher), dystrophin (MA1-26837, Invitrogen), and laminin (L9393, Sigma-Aldrich) were incubated overnight at 4°C. Samples were washed three times with PBS and Alexa Fluor secondary antibodies (Invitrogen) were incubated for 1h at RT. After washing three times with PBS, a cover slide was placed with mounting media with or without DAPI counterstaining. All slides were imaged under an inverted microscope (Leica DMi8) and analyzed in a blinded manner by ImageJ (NIH) and Las X software (Leica).

Plethysmography

Respiratory function was assessed in new-born mouse pups (postnatal day 1, P1) using a custom-designed plethysmograph as described (*4*). Prior to measurements, barometric chambers were normalized using a Hamilton syringe. Animals were then placed in the experimental barometric chamber and equilibrated for 5 minutes on a heated pad. Then, the chamber was sealed and differential pressure between the experimental and reference chambers measured using a pressure transducer. Approximately 50 consecutive breaths were analyzed using LabChart 7.0 software (AD Instruments) to measure respiration rate, tidal volume, and breathing regularity.

Whole body X-ray scanning

Newborn WT and *Mybpc1*$^{-/-}$ mice were sacrificed within 24 hrs after birth and fixed in 4%PFA. CT scans of ~27μm were acquired on fixed mice using a Siemans Inveon PET/SPECT/CT and sections combined for 3D rendering of bone structure.

Treadmill running and grip strength tests

As previously described (*3*), a graded running test was used to evaluate a running capacity. After three consecutive days of practice running at the speed of 4 to 10m/min for 10 min, maximum running capacity on the treadmill (Omnitech Electronics, Columbus, OH, USA) was measured with increment speed and angle every 1-2 min. Total running distance and time before exhaustion was recorded to define the maximum running capacity.

Forelimb grip strength was measured three times by pulling the mouse against a grip strength meter (1027SM, Columbus Instruments, Columbus, OH, USA). The peak value was normalized for individual body weight and compared between groups.

*In vivo* hindlimb muscle function

To assess muscle strength *in vivo*, we conducted measurements of isometric peak tetanic torque ($P_o$) of the plantarflexor muscles, following a previously established protocol (*3*). After anesthetizing with 2% vaporized isoflurane, the right knee of each mouse was secured at a 90-

degree angle, with the foot affixed to a 2 cm lever on a dual-mode servomotor system (300C-LR: Aurora Scientific, Aurora, ON, Canada). $P_o$ was recorded in response to evoked contractions, achieved by stimulating the tibia nerve via two needle electrodes connected to an electrical stimulator (701C: Aurora Scientific, Aurora, ON, Canada). Repetitive stimulations lasting 0.2 msec were administered at a frequency of 50 to 200Hz, for 350 msec, every two minutes.

*Ex vivo* intact muscle function test

Intact EDL and SOL muscles were isolated and placed in a tissue bath (25 ml) that was perfused with oxygenated (95% O2 and 5% CO2) Krebs-Henseleit buffer (pH 7.40 at 36$^{o}$C). One tendon was fixed, while the other was attached to a dual-mode lever arm using silk suture. The muscle was left to equilibrate for 10 minutes before stimulation. Electric field stimulation was applied using 0.2 msec pulses at 500 mA delivered via a pair of platinum plate electrodes placed parallel to the muscle, using an electrical stimulator (701C: Aurora Scientific, Aurora, ON, Canada). The muscle was adjusted to the optimal length ($L_o$) for generating peak isometric twitch force ($P_t$). Peak isometric tetanic force ($P_o$) was calculated using tetani of 350 msec duration and frequencies of 12.5 to 200 Hz applied every two minutes. Fatigue resistance was assessed by inducing repeated isometric tetanic contractions at a frequency of 150Hz, with each contraction delivered at 10-second intervals for a total of 50 contractions. The percentage loss of force generation after every 10 contractions was calculated and graphed relative to the initial force. Muscle mass and length were measured after all the tests. Physiological muscle cross-sectional area (CSA) was calculated as [Muscle weight (g)]/ [fiber density (1.06mg/mm$^3$) x optimal fiber length (mm)]. Optimal fiber length was determined by dividing $L_o$ by muscle length to fiber length ratios, 0.44 for EDL and 0.71 for Soleus (2). Specific force ($SP_o$) was calculated by normalizing $P_o$ by CSA ($P_o$/CSA) (N/cm$^2$). In vivo and ex vivo muscle function data were collected using a commercial muscle lever system (300C-LR: Aurora Scientific, Aurora, ON, Canada), digitalized using Dynamic Muscle Control (DMC v5.5), and analyzed using Dynamic Muscle Analysis (DMA v5.3) (Aurora Scientific, Aurora, ON, Canada).

*In vitro* skinned fiber test

Dissected diaphragm and soleus muscles were incubated in a glycerinated relaxing solution (7 mM EGTA, 100 mM BES, 0.017 mM CaCl2, 5.49 mM MgCl2, 5 mM DTT, 15 mM creatine phosphate, 4.66 mM ATP, 55.7 mM K-propionate, pH 7.0 and 50% (v/v) glycerol) overnight at 4$^{o}$C on a rotating shaker. The relaxing solution was changed twice a day for two days to remove muscle membrane debris. The skinned samples were stored at -20$^{o}$C with fresh glycerinated relaxing solution until use.  Skinned muscle samples were dissected into single or few fibers to measure the isometric force generation, and aluminum T-clips were attached to both sides. The fiber was transferred to Aurora Scientific lever system (405A, Aurora, ON, Canada) equipped with a force transducer and length controller. Sarcomere length was adjusted to ~2.5 µm, and fiber diameter was determined by Aurora Scientific Video Sarcomere Length software. Isometric force was measured at different concentrations of $Ca^{++}$ ranging from pCa 9.0 to pCa 4.5 in activation buffer (7 mM EGTA, 100 mM BES, 7.01 mM CaCl2, 5.29 mM MgCl2, 5 mM DTT, 15 mM creatine phosphate, 4.72 mM ATP, and 55.7 mM K-propionate at pH 7.0). The fiber was washed with a relaxing solution for two minutes after each contraction and force was normalized by fiber cross-sectional area (CSA). To measure the rate of cross-bridge cycling in skinned fibers at pCa 4.5, force redevelopment rate (ktr) was measured by the slack and re-stretch test as previously described (*5*). Acquired results were analyzed using a single exponential equation, and the half-life of tension redevelopment was used to calculate Ktr (s-1) after linear transformation. All of the data were collected using the Real-time Muscle Acquisition and Analysis System (600A: Aurora Scientific, Aurora, ON, Canada).

RNAseq

Diaphragms from P1 and adult soleus muscles were homogenized in PureZOL (Bio-Rad, #7326890) or RLT lysis buffer (QIAGEN, 79216), respectively. Total RNA was extracted using a RNeasy Mini Kit (Qiagen, 74104) with additional in-column DNase1 treatment (79254, Qiagen). The quality and quantity of the isolated RNA was measured by the Agilent 2100 Bioanalyzer. mRNA was purified from total RNA using oligo(dT)-attached magnetic beads and fragmented into small pieces. cDNA was synthesized with random hexamers followed by second-strand synthesis and purification. A cDNA library was created by PCR enrichment and validated on the Agilent Technologies 2100 bioanalyzer. Libraries were pooled and sequenced by 150bp paired-end run using an Illumina Hiseq2500 for diaphragm and 100bp paired-end run using an DNBSEQ-G400 for soleus with at least 40 million reads per sample.
Base calling was performed using Illumina CASAVA (v1.4), and the quality of sequencing reads was checked using FASTQC (v0.11.7). Reads were aligned against mm10 mouse genome with HISAT2 (v2.0.5) (14). Raw gene counts were calculated using feature Counts (v1.5.2) (15) and normalized using edgeR's TMM (trimmed mean of M values) method. Differentially expressed genes (DEGs) were detected using limma/voom. Genes exceeding the moderate cutoff set (log FC > 1.5 and p < 0.001 or log FC > 1.5 and FDR<0.05 ) were considered differentially expressed. These DEGs were used as queries to EnrichR and ClueGO to retrieve enriched pathways and/or gene ontology terms. The GSEA analysis was also performed by gene-set permutation.

Muscle X-ray diffraction

Small-angle X-ray diffraction patterns on soleus muscles were collected using the BioCAT beamline 18ID at the Advanced Photon Source, Argonne National Laboratory as described previously (*6*). Briefly, intact soleus muscle was mounted vertically in the Aurora Scientific lever system (300C-LR, Aurora, ON, Canada) and immersed in oxygenated Krebs-Henseleit buffer (pH 7.40) at room temperature (24 $^{o}$C). Isometric tetanic force was generated by electrical stimulation at 150Hz for 1.5sec with 0.2 m pulse after 1.0 s initial delay. The X-ray beam energy was set to 12KeV (0.1033 nm wavelength) with an incident flux of ~$10^{13}$ photons per second. The X-ray beam dimensions were about 250 µm x 250 µm at the sample and approximately 150 x 30 µm at the detector (Pilatus 3 1M, Dectris Inc., Baden, Switzerland). The specimen to detector distance was approximately 3 meters. During an isometric tetanic muscle contraction, data was collected continuously at 50 frames per second with a 10 msec X-ray exposure and 10 msec per frame readout. To minimize radiation damage, the sample was oscillated vertically and horizontally at a velocity of 10 mm/s, which corresponded to the X-ray beam dimensions, to avoid overlapping X-ray exposures on the sample surface during data collection.
Raw X-ray images during resting and plateau region were averaged using FIT2D software. Interfilament lattice spacings and 1,1 to 1,0 equatorial reflection intensity ratios were calculated using the Equator module in MuscleX software (*7*) as previously described (*8*). The diffraction images were quadrant folded and had the diffuse background images subtracted using Quadrant Fold program in MuscleX software for subsequent analysis. The meridional spacing and intensities and the intensities of myosin-based layer line were measured using Projection Trace module in MuscleX software as described previously (*9*).

Transmission electron microscopy

Whole soleus muscles were dissected and secured to wooden sticks at physiological length in relaxing buffer (5% dextrose, 0.15% sucrose and 100mM KCl in PBS). After incubating in 1% paraformaldehyde and 2% glutaraldehyde in relaxing buffer for 1hr on ice, samples were detached and placed in 1% paraformaldehyde and 2% glutaraldehyde in 0.1M cacodylate buffer, pH 7.2 overnight at 4$^{o}$C. Tissues were postfixed in 1% osmium tetroxide in 0.15M sodium cacodylate buffer, processed through a series of alcohols, infiltrated, and embedded in LX-112 resin. After polymerization at 60 degrees for three days, ultrathin sections (120nm) were cut using a Leica EM UC7 ultramicrotome and counterstained in 2% aqueous uranyl acetate and Reynold's lead citrate. Images were taken with a transmission electron microscope (Hitachi H-7650) equipped with a digital camera (Biosprint 16) and analyzed using ImageJ (NIH).

Statistical Analysis

All data is presented as mean ± SE. For group comparisons, we used Student's t-test or one way ANOVA with Tukey's post-hoc test using the GraphPad Prism 7.04 software. Survival curves of new-born pups were analyzed by Mantel-Cox test. Statistical significance was defined as a P-value less than 0.05.
